# A druggable chromatin vulnerability in suppressive neutrophils restores antitumor immunity

**DOI:** 10.64898/2026.09.26.754703

**Authors:** Fan Yang, Yi Bao, Yuanyuan Qiao, Gabriel Cruz, Eleanor Young, Rahul Mannan, Yuping Zhang, Matt Goodrich, Yizhi Cao, Jie Luo, Radha Paturu, Somnath Mahapatra, Jean Ching-Yi Tien, Royce Xu, Mikoto Kobayashi, Yihan Liu, Andrej Coleski, Fengyun Su, Xuhong Cao, Stephanie J. Miner, Linda Vatan, Ranga Sudharshan, Arvind Rao, Susanta Samajdar, Chandrasekhar Abbineni, Murali Ramachandra, Jessica C. Piel, Martin Hentemann, Ilona Kryczek, Weiping Zou, Lanbo Xiao, Arul M. Chinnaiyan

## Abstract

Many cancers evade immunity by converting neutrophils into polymorphonuclear myeloid_-_derived suppressor cells (PMN-MDSCs), but the chromatin mechanisms that maintain this pathological state remain unclear. Here we show that tumor-induced suppressive neutrophils depend on mSWI/SNF chromatin remodeling. Two orally bioavailable mSWI/SNF ATPase antagonists, a degrader and a catalytic inhibitor, produced immune-dependent tumor control across syngeneic models, including tumors resistant to PD-1 blockade, and enhanced checkpoint therapy without overt toxicity. Treatment rapidly depleted intratumoral PMN-MDSCs, restored functional CD8^+^ T cells, and blocked tumor explant-induced expansion and suppressive activity of mouse and human PMN-MDSCs. mSWI/SNF antagonism collapsed chromatin accessibility and PU.1/C/EBPβ occupancy at regulatory elements that sustain PMN-MDSC identity. Thus, suppressive neutrophils harbor a druggable chromatin vulnerability that can be exploited to restore cancer immunity.

## Introduction

Many cancers evade immune destruction by remodeling the tumor microenvironment rather than by disabling T cells alone [1-3]. A major component of this immune resistance is the accumulation of polymorphonuclear myeloid-derived suppressor cells (PMN-MDSCs), also known as pathologically activated suppressive neutrophils [4, 5]. These cells arise during tumor_-_induced emergency granulopoiesis, accumulate in many solid tumors, suppress CD8^+^ T cell function, and are associated with resistance to immune checkpoint blockade [6, 7]. Although substantial efforts have focused on blocking PMN-MDSC trafficking or effector pathways, these approaches have produced limited and often incomplete activity [8, 9]. A central unresolved question is what molecular machinery maintains the pathological neutrophil state once it is induced by cancer.

Cellular identity is enforced by chromatin regulatory programs that determine which transcriptional circuits remain accessible and active [10]. The mSWI/SNF family of ATP_-_dependent chromatin remodeling complexes shapes enhancer and promoter accessibility across development, differentiation, and cancer [11-14]. In oncology, mSWI/SNF complexes have been studied largely as tumor cell-intrinsic dependencies, particularly in transcription factor-driven malignancies, leading to the development of selective, orally bioavailable mSWI/SNF ATPase antagonists [13, 15]. mSWI/SNF complexes also regulate immune-cell differentiation and function, including T cell fate and exhaustion programs [16-18], but whether systemic mSWI/SNF antagonism can dismantle suppressive immune states within the tumor microenvironment has remained unclear.

Here we identify mSWI/SNF chromatin remodeling as a druggable dependency of tumor_-_induced suppressive neutrophils. Using two mechanistically distinct mSWI/SNF ATPase antagonists, the degrader AU-24118 [19] and the catalytic inhibitor FHD-286 [20], we show that systemic mSWI/SNF antagonism elicits immune-dependent tumor control across syngeneic cancer models, including tumors resistant to PD-1 blockade, and enhances checkpoint therapy without overt toxicity. Unbiased immune profiling, genetic perturbation, and depletion studies identify PMN-MDSCs as the primary immune target and CD8^+^ T cells as the required effector population. mSWI/SNF antagonism blocks tumor-induced expansion and suppressive activity of mouse and human PMN-MDSCs and collapses PU.1_-_and C/EBPβ-associated chromatin accessibility at regulatory elements that maintain the PMN-MDSC program. These findings reveal a druggable chromatin vulnerability in suppressive neutrophils and establish mSWI/SNF antagonism as a strategy to restore antitumor immunity.

## Results

### Systemic mSWI/SNF antagonism restores immune control of checkpoint-resistant tumors

To evaluate the therapeutic efficacy of systemic administration of mSWI/SNF antagonists, we tested AU-24118 and FHD-286 across a broad panel of syngeneic tumor models encompassing both immune checkpoint blockade (ICB)-sensitive and -resistant settings. In ICB-sensitive models, including A20 lymphoma and MC38 colon adenocarcinoma, treatment with AU-24118 or FHD-286 elicited robust antitumor activity comparable to ICB (**Figure 1A**). Importantly, in ICB_-_insensitive models, including B16-F10 melanoma, CT26 colon adenocarcinoma, and 4T1 triple_-_negative breast cancer, mSWI/SNF antagonism demonstrated superior antitumor efficacy relative to ICB in wild-type mice (**Figure 1A**). Strikingly, the therapeutic activity of both compounds was strictly dependent on an intact immune microenvironment, as neither agent reduced tumor burden in these models, in immune-deficient NOD.Cg-*Prkdc^scid^ Il2rg^tm1Wjl^*/SzJ (NSG) mice (**Figure 1B**). To confirm on-target activity, we verified efficient depletion of the mSWI/SNF ATPase subunits BRG1 and BRM following AU-24118 treatment by immunofluorescence analysis (**Figure S1A**). Together, these findings indicate that systemic mSWI/SNF antagonism drives potent, immune-dependent antitumor responses across multiple tumor contexts, including models refractory to ICB.

**Figure 1.**
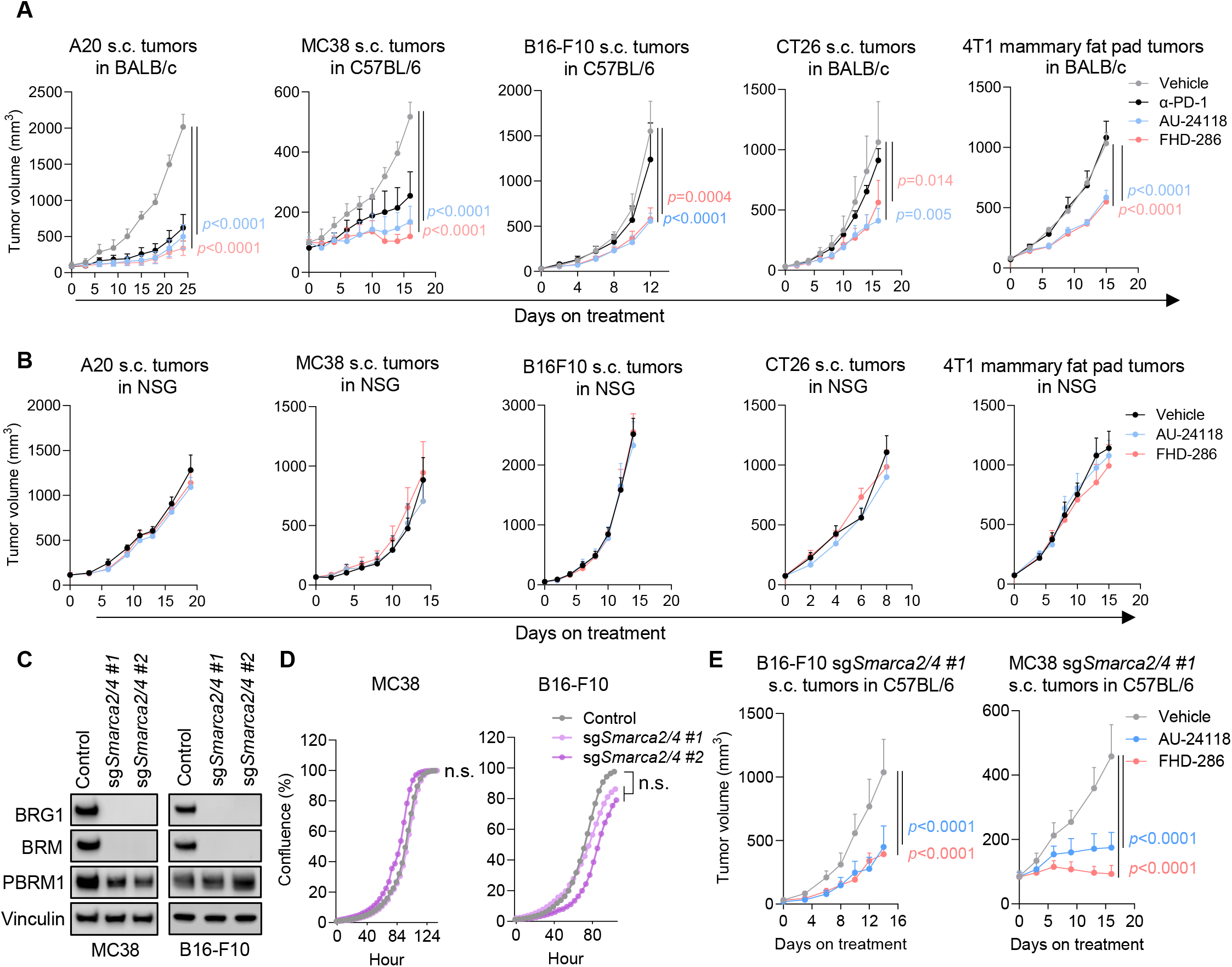
mSWI/SNF antagonists display immune-dependent antitumor activity. A. Temporal changes in tumor volume of the indicated models in immunocompetent syngeneic mice treated with AU-24118 (7.5 mg/kg), FHD-286 (1.5 mg/kg), anti-PD-1 (100 μg per mouse for MC38 tumor models and 200 μg per mouse for other models) and vehicle. s.c.: subcutaneous. α-PD-1: anti-PD-1. *n* = 5–6 mice per group. B. Temporal changes in tumor volume of the indicated models in NOD.Cg-*Prkdc^scid^ Il2rg^tm1Wjl^*/SzJ (NSG) mice treated with AU-24118, FHD-286, and vehicle. *n* = 6–7 mice per group. C. Immunoblots showing levels of the indicated proteins in MC38 and B16-F10 cells that received a nontargeting CRISPR guide sequence (control) or distinct sgRNAs depleting *Smarca2* and *Smarca4*. Vinculin is used as a loading control. D. Cell proliferation measurement of cells in **C** (*n* = 3 technical replicates). E. Temporal changes in tumor volume of tumors derived from cells in **C**, in syngeneic mice treated with AU-24118, FHD-286, and vehicle. *n* = 8–10 mice per group. All data are presented as mean ± SEM and were statistically assessed using two-way analysis of variance. Bonferroni’s correction is applied for multiple comparisons in **A** and **E**. All data are representative of two independent experiments.

To further confirm the immune dependence of the observed therapeutic efficacy, we generated tumor cells with complete knockout of *Smarca2* and *Smarca4* (*Smarca2/4*), which encode BRM and BRG1, respectively, using independent CRISPR guide RNA sequences (sg*Smarca2/4* #1 and sg*Smarca2/4* #2) (**Figure 1C**). *Smarca2/4* loss did not impair tumor-cell proliferation in vitro (**Figure 1D**), indicating that these models are not intrinsically dependent on mSWI/SNF ATPase activity for growth. Importantly, pharmacologic mSWI/SNF antagonism remained effective in suppressing the growth of *Smarca2/4*-deficient tumors in vivo, further demonstrating that the antitumor response is mediated through the immune compartment rather than tumor cell intrinsic vulnerabilities (**Figure 1E**).

We next evaluated the therapeutic efficacy of combining mSWI/SNF antagonism with ICB. In the A20 model, monotherapy with AU-24118 or FHD-286 achieved complete response (CR) rates of 14% (1/7 CR) and 25% (2/8 CR), respectively (**Figure 2A**), leading to prolonged tumor-free survival in the treated animals through the end of the study (**Figure 2B**). Notably, combination treatment with anti-PD-1 and either AU-24118 or FHD-286 resulted in the highest CR rates of 50% (4/8 CR) and 75% (6/8 CR), respectively, markedly exceeding those achieved with either single agent alone (**Figure 2A**). Consistent with these findings, combination therapy led to a substantial extension of tumor-free survival compared with anti-PD-1 monotherapy (**Figure 2B**). Similar combinational effects were observed in additional syngeneic models, including MC38 (**Figure 2D** and **Figure S1B**) and, notably, the ICB-insensitive CT26 model (**Figure 2C**). Importantly, both monotherapy and combination therapy were well tolerated, with no evidence of systemic toxicity or body weight loss (**Figure S1C**). This is in strong agreement with prior studies of AU-24118 [21, 22] and FHD-286 [23], which likewise reported favorable tolerability.

**Figure 2.**
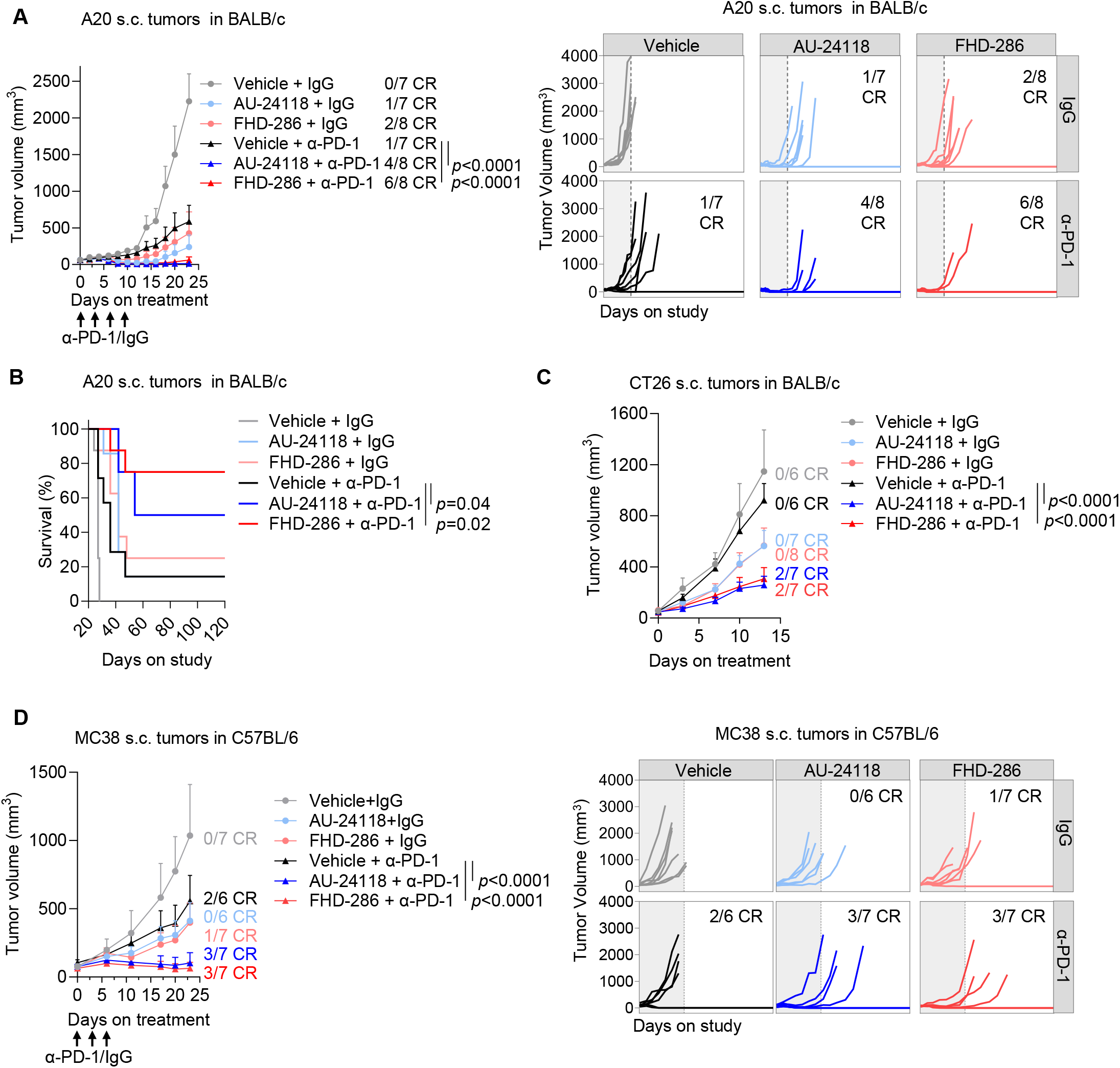
mSWI/SNF antagonists synergize with PD-1 blockade across multiple syngeneic mouse models. A. Temporal changes in tumor volume of the indicated tumors in syngeneic mice treated with vehicle, AU-24118, or FHD-286 in combination with IgG or anti-PD-1 (*n* = 7–8 mice per group). The gray regions represent the treatment period. CR: complete response. B. Survival analysis of mice treated in A. C. Temporal changes in tumor volume of CT26 s.c. tumors in syngeneic mice treated with vehicle, AU-24118, or FHD-286 in combination with IgG or anti-PD-1 (*n* = 6–8 mice per group). D. Temporal changes in tumor volume of MC38 subcutaneous (s.c.) tumors in syngeneic mice treated with vehicle, AU-24118, or FHD-286 in combination with IgG or anti-PD-1(*n* = 6–7 mice per group). All data are presented as mean ± SEM and were statistically assessed using two-way analysis of variance, except for B which used the log-rank test. Bonferroni’s correction is applied for multiple comparisons in A, B and D. All data are representative of two independent experiments.

### mSWI/SNF antagonism depletes suppressive neutrophils and restores CD8^+^ T cell immunity

To define the immune mediators required for immune-dependent therapeutic efficacy, we performed single-cell RNA sequencing (scRNA-seq) of CD45_⁺_ tumor-infiltrating immune cells isolated from MC38 tumors following mSWI/SNF antagonism (**Figure 3A** and **Figure S2A**). This unbiased analysis revealed that both AU-24118 and FHD-286 elicited profound remodeling of the tumor microenvironment, marked by a pronounced depletion of polymorphonuclear myeloid_-_derived suppressor cells (PMN-MDSCs, also known as pathologically activated neutrophils) (**Figure 3A–B**), which suppress immune effector cells in tumors [4, 24, 25]. Consistently, a concomitant expansion of immune effector cells, particularly CD8_⁺_ T cells, was also observed (**Figure 3A–B**). Among all myeloid subsets, the reduction in PMN-MDSCs was the most substantial (**Figure 3B**). These findings were independently validated by flow cytometry (**Figure S2B**), which confirmed a marked decrease in PMN-MDSC abundance alongside robust increases in both total and functional CD8_⁺_ T cells expressing Ki67, TNFα, IFN-γ, and granzyme B (**Figure 3C** and **Figure S3A**). In contrast, other immune populations, such as monocytic MDSCs, regulatory T cells, B cells, dendritic cells, and macrophages, remained largely unchanged (**Figure S3A**), highlighting that systemic mSWI/SNF antagonism selectively depletes PMN-MDSCs within the tumor microenvironment at the therapeutically effective doses. Temporal analysis revealed that PMN-MDSC depletion occurred as early as 36 hours following treatment initiation, preceding detectable changes in CD8_⁺_ T cell or NK cell populations (**Figure 3D**). These kinetics indicate that direct modulation of PMN-MDSCs represents a primary event driving subsequent immune remodeling. Parallel analyses across multiple syngeneic models including 4T1, B16-F10, CT26, and A20 demonstrated that mSWI/SNF antagonism consistently suppressed intratumoral PMN-MDSC accumulation while promoting the expansion of functional CD8_⁺_ T cells (**Figure 3E** and **Figure S3B–E**). Notably, an increase in NK cells was observed only in the MC38 model (**Figure 3C**) following PMN-MDSC reduction, but not in the other models examined (**Figure S3B–C**), suggesting that CD8_⁺_ T cells constitute the principal immune effector population restrained by PMN-MDSCs across most tumor contexts tested [4, 24, 25].

**Figure 3.**
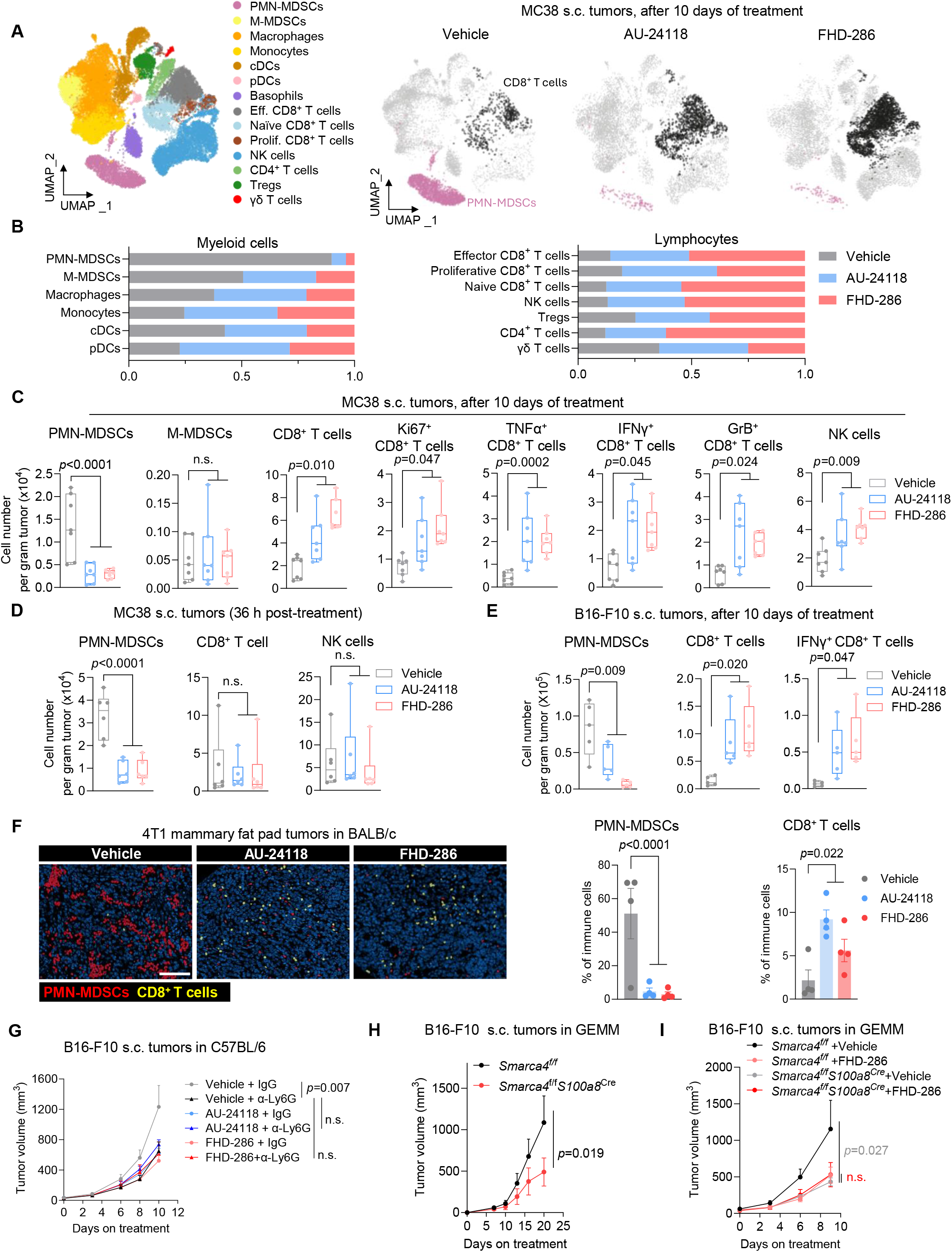
mSWI/SNF antagonism depletes PMN-MDSCs while increasing functional CD8^+^ T cells in tumors. A. scRNA-seq analysis of CD45^+^ leukocytes sorted from MC38 subcutaneous (s.c.) tumors in mice treated with vehicle, AU-24118, or FHD-286. Tumors from five mice per group were combined. Merged uniform manifold approximation and projection (UMAP) plot (left) and density plots for each treatment group (right) are shown. PMN-MDSCs: polymorphonuclear myeloid-derived suppressor cells; M-MDSCs: monocytic myeloid_-_derived suppressor cells; cDCs: conventional dendritic cells; pDCs: plasmacytoid dendritic cells; Eff.: Effector; Prolif.: Proliferative; NK: natural killer; Tregs: regulatory T cells. B. Proportion of immune subpopulations in each group from A. C. Flow cytometric analysis of absolute numbers of the indicated immune cells in MC38 s.c. tumors from mice treated with vehicle (gray), AU-24118 (blue), and FHD-286 (pink) for 10 days (*n* = 7 mice per group). n.s.: not significant; TNF: tumor necrosis factor; IFN: interferon; GrB: granzyme B. D. Flow cytometric analysis of absolute numbers of PMN-MDSCs, CD8_⁺_ T cells, and NK cells in MC38 s.c. tumors at a short-term time point (36 hour) following treatment with vehicle (gray), AU-24118 (blue), and FHD-286 (pink) (*n* = 6 mice per group). E. Flow cytometric analysis of absolute numbers of the indicated immune cells in B16-F10 s.c. tumors from mice treated with vehicle (gray), AU-24118 (blue), and FHD-286 (pink) for ten days (*n* = 5 mice per group). F. Representative Xenium In Situ spatial plots from 4T1 mammary fat pad tumors treated with vehicle, AU-24118, and FHD-286 (Left). PMN-MDSCs are shown in red, and CD8^+^ T cells are shown in yellow. Scale bar is 200 µm. Quantification (right) of PMN-MDSCs and CD8^+^ T cells identified in Xenium In Situ. G. Tumor volume changes of B16-F10 s.c. tumors treated with vehicle, AU-24118, or FHD_-_286 in combination with IgG or anti-Ly6G (*n* = 8 mice per group). α-Ly6G: anti-Ly6G. H. Tumor volume changes of B16-F10 s.c. tumors in *Smarca4^f/f^* or *Smarca4^f/f^S100a8^cre^* mice (*n* = 7–8 mice per group). I. Tumor volume changes of B16-F10 s.c. tumors in *Smarca4^f/f^* or *Smarca4^f/f^S100a8^cre^* mice treated with vehicle or FHD-286 (*n* = 8 mice per group). All data are presented as box and whisker plots except B (bar plot), G, H, and I (mean ± SEM), and were statistically assessed using two-way analysis of variance, except B. Bonferroni’s correction is applied for multiple comparisons in G and I. All data except A, B, and F are representative of two independent experiments.

To validate the immune landscapes defined by scRNA-seq and flow cytometry, we further performed high-resolution in situ spatial transcriptomic profiling using the Xenium In Situ platform (10x Genomics) on tumors treated with or without mSWI/SNF antagonism. The subcellular spatial resolution afforded by this platform enabled delineation of the cellular composition and spatial organization of the tumor microenvironment in situ. Using the Xenium Prime 5K Mouse Pan Tissue & Pathways panel supplemented with a custom 100-gene add-on panel (**Supplementary Table 1**), we resolved cellular clusters including myeloid, T/NK-cell, B cell, stromal, and endothelial compartments (**Figure S4A–C**). Notably, this analysis recapitulated the inverse regional association between PMN-MDSC accumulation and CD8^+^ T cell infiltration (**Figure 3F**).

To establish the functional requirement of these immune subsets, we performed antibody_-_mediated depletion studies. Depletion of PMN-MDSCs using anti-Ly6G antibodies [26, 27], which target Ly6G-expressing mature granulocytic cells including PMN-MDSCs, inhibited tumor growth but failed to confer additional benefit when combined with mSWI/SNF antagonists, consistent with a shared mechanistic axis between the antibody and these agents (**Figure 3G** and **Figure S5A**). This finding further indicates that PMN-MDSCs represent the principal pharmacological target of mSWI/SNF antagonists. In contrast, depletion of CD8_⁺_ T cells completely abrogated the anti-tumor efficacy of both AU-24118 and FHD-286, establishing CD8_⁺_ T cells as the indispensable effector population mediating tumor control (**Figure S5B**). Collectively, these data indicate that mSWI/SNF antagonism depletes intratumoral PMN-MDSCs while increasing immune-effector CD8_⁺_ T cells, thereby promoting antitumor immunity and suppressing tumor growth.

To further define the role of mSWI/SNF in PMN-MDSCs, we used a genetic approach to specifically deplete the mSWI/SNF ATPase subunit. To this end, we crossed transgenic *Smarca4*^f/f^ mice [28] with *S100a8-Cre* mice [29, 30] to conditionally knockout *Smarca4* in PMN_-_MDSCs. Ablation of *Smarca4* in PMN-MDSCs was sufficient to inhibit tumor growth (**Figure 3H**), accompanied by a reduction in intratumoral PMN-MDSC abundance and an increase in CD8_⁺_ T cell infiltration, supporting a *Smarca4*-dependent requirement for the accumulation of tumor_-_associated PMN-MDSCs (**Figure S5C**). This finding is consistent with the notion that BRG1 (encoded by *Smarca4*) plays a dominant and often non-redundant role in mSWI/SNF function relative to BRM (encoded by *Smarca2*) [13, 31, 32]. Importantly, mSWI/SNF antagonism did not confer additional anti-tumor benefit in *Smarca4*^f/f^*S100a8^Cre^* mice (**Figure 3I**), suggesting that loss of BRG1 accounts for much of the mSWI/SNF dependency observed in vivo.

Other agents have also been reported to deplete PMN-MDSCs in tumors. Among these, tasquinimod has advanced to Phase III clinical trials in metastatic castration-resistant prostate cancer [8, 30, 33]. Notably, mSWI/SNF antagonists achieved superior tumor control and more profound PMN-MDSC depletion compared with tasquinimod (**Figure S5D**), suggesting improved therapeutic potential relative to tasquinimod.

### mSWI/SNF antagonism blocks tumor-induced suppressive-neutrophil expansion and function

To elucidate the molecular programs underlying the consistent depletion of PMN-MDSCs, we performed transcriptional profiling of tumor-associated PMN-MDSCs isolated following mSWI/SNF antagonism. Gene ontology analysis revealed that mSWI/SNF antagonism profoundly suppressed pathways central to PMN-MDSC function, including neutrophil activation, activation involved in immune response, degranulation, and extravasation (**Figure 4A and Figure S6A**). This transcriptional reprogramming was marked by robust downregulation of a core set of PMN-MDSC-associated functional genes, such as *Ngp*, *Camp*, *Ltf*, *Cd177*, *S100a8*, and *S100a9* (**Figure 4B**) [30, 34, 35]. These changes were independently validated in vitro, where mSWI/SNF antagonism induced dose-dependent transcriptional repression of these genes (**Figure 4C**). Notably, these changes occurred within 24 hours, consistent with the rapid in vivo depletion of PMN-MDSCs observed within 36 hours (**Figure 3D**). Guided by the RNA_-_seq analysis, we incorporated a custom 100-gene add-on panel into the Xenium Prime 5K Mouse Pan Tissue & Pathways panel (**Supplementary Table 1**). This enabled in situ assessment of the PMN-MDSC program and revealed concordant downregulation of the PMN_-_MDSC-associated functional gene signature in mSWI/SNF antagonist-treated tumors relative to vehicle-treated controls, consistent with the RNA-seq findings (**Figure 4D**). To determine whether these transcriptional changes were reflected at the protein and tissue-architectural levels, we performed multiplex immunofluorescence. AU-24118 reduced BRG1 and BRM expression in tumor-infiltrating PMN-MDSCs, confirming on-target mSWI/SNF ATPase suppression in this compartment (**Figure S6B**). Consistent with RNA-seq and Xenium In Situ profiling, mSWI/SNF antagonism diminished S100A9 and CD177 protein expression in residual S100A9^+^ PMN-MDSCs (**Figure S6C**). Collectively, these data show that mSWI/SNF antagonism attenuates expression of PMN-MDSC-associated functional genes.

**Figure 4.**
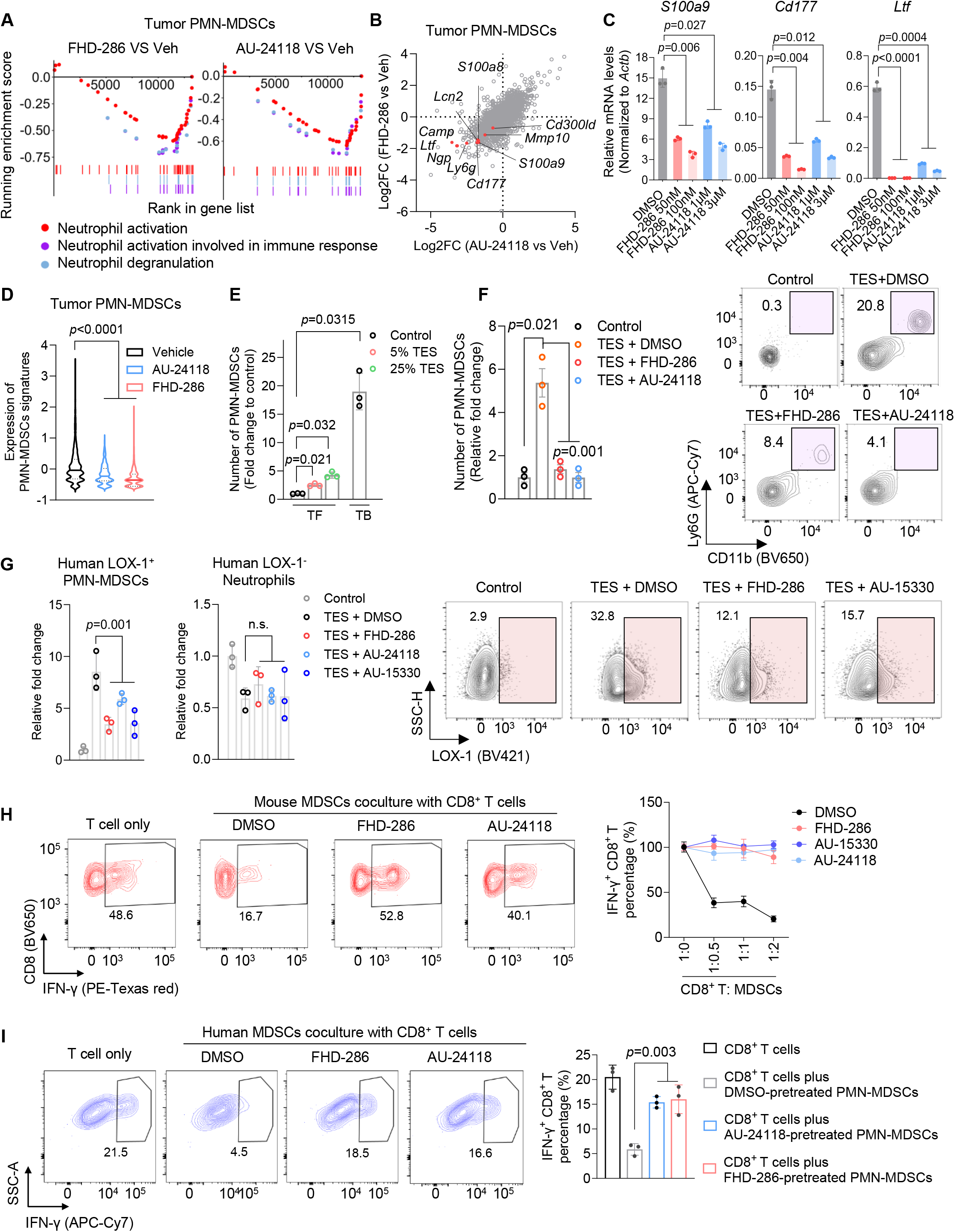
mSWI/SNF antagonism attenuates tumor-induced PMN-MDSC expansion and function. A. Gene Ontology pathways enriched in bulk RNA-sequencing profiles of PMN-MDSCs from tumors in mice treated with FHD-286 or AU-24118 compared with vehicle (Veh.). PMN-MDSCs were isolated from five mice per group. B. Scatter plot assessed by RNA-sequencing in A, visualizing the overall transcriptomic alterations. PMN-MDSC-associated functional genes are highlighted in red. C. Reverse transcription quantitative PCR assessing expression of *S100a9*, *Cd177*, and *Ltf* in PMN-MDSCs treated with DMSO, FHD-286, or AU-24118 with different concentrations in vitro for 24 hours. D. Xenium In Situ analysis showing the expression of PMN-MDSC signature genes (*Ltf, Camp, Ly6g, Ngp, Cxcr2, Cd177, Wfdc21, Chil3, Hdc,* and *Cd84*) in 4T1 mammary fat pad tumors following treatment with vehicle, AU-24118, or FHD-286. E. Flow cytometric quantification of absolute PMN-MDSC numbers in splenocytes isolated from tumor-free (TF) mice cultured in medium alone or supplemented with 5% or 25% tumor explant supernatant (TES), as well as in splenocytes isolated from tumor-bearing (TB) mice. F. Left: Flow cytometric quantification of absolute PMN-MDSC numbers in splenocytes isolated from TF mice cultured in medium alone or supplemented with 25% TES, with 24-hour in vitro treatment by DMSO, FHD-286 (100 nM), or AU-24118 (3 µM). Right: Representative contour plots showing the proportional change of PMN-MDSCs treated in left. G. Left: Flow cytometric quantification of absolute numbers of human LOX-1_⁺_ PMN-MDSCs and LOX-1_⁻_ neutrophils in cells isolated from patients’ whole blood. Cells were cultured for 24 hours in medium alone or supplemented with 25% patient tumor-derived TES and treated with FHD-286 (100 nM), AU-24118 (3 µM), or AU-15330 (3 µM). Cells and TES were obtained from three individual patients. Right: Representative contour plots showing the corresponding changes in the proportion of LOX-1_⁺_ PMN-MDSCs. H. Representative flow cytometry plots (left) and quantitative analysis (right) showing the relative fold changes of percentage of IFN-γ_⁺_ CD8_⁺_ T cells after coculture with mouse PMN-MDSCs that had been pretreated for 24 hours with DMSO, FHD-286 (100 nM), AU-24118 (3 µM), or AU-15330 (3 µM), at the indicated ratios for 72 hours. I. Representative flow cytometry plots (left) and quantitative analysis (right) showing the relative fold changes of percentage of IFN-γ_⁺_ CD8_⁺_ T cells after coculture with human PMN-MDSCs that had been pretreated for 24 hours with DMSO, FHD-286 (100 nM), or AU-24118 (3 µM) at the indicated ratios for 72 hours. Data are presented as box and whisker plots, except in A, B, and H (right, mean ± SEM). Significance in C-G and I was determined by two-way analysis of variance. Bonferroni’s correction is applied for multiple comparisons in C, E and F. All data are representative of two independent experiments.

Tumor progression is known to induce “emergency granulopoiesis”, a systemic adaptation in which hematopoietic organs preferentially generate immature myeloid cells at the expense of terminally differentiated effector populations [36-38]. Sustained exposure to tumor-derived inflammatory cytokines and growth factors during this process drives the functional reprogramming of neutrophils toward an immunosuppressive PMN-MDSC phenotype [37-39]. Consistent with this paradigm, splenocytes isolated from tumor-bearing (TB) mice exhibited a marked expansion of PMN-MDSCs (CD11b_⁺_ Ly6G_⁺_) compared with those from tumor-free (TF) controls (**Figure 4E and Figure S7A**). We further demonstrated that tumor explant supernatants (TES) derived from various tumor models were sufficient to recapitulate this pathological differentiation in vitro, inducing a significant expansion of PMN-MDSCs (**Figure 4E** and **Figure S7B**). These findings confirm that soluble tumor-derived factors are the principal drivers of MDSCs expansion [4, 39]. Importantly, mSWI/SNF antagonism effectively abrogated TES-induced PMN-MDSC expansion (**Figure 4F**) and suppressed TES-induced expression of PMN-MDSCs associated genes (**Figure S7C**). These observations were further validated in human systems, where mSWI/SNF antagonism selectively suppressed TES-induced expansion of human PMN-MDSCs (LOX-1_⁺_) [40] (**Figure 4G and Figure S7D**), while sparing homeostatic human neutrophils (LOX-1_⁻_). This is consistent with in vivo immune profiling, which revealed that systemic mSWI/SNF antagonism did not alter the abundance of intratumoral myeloid populations other than PMN-MDSCs (**Figure 3C** and **Figure S3A**). Collectively, these data identify the mSWI/SNF ATPase activity as an essential molecular requirement for the transcriptional programs that enable tumor-driven PMN-MDSC expansion across species.

The suppression of neutrophil activation and extravasation programs identified by bulk RNA sequencing (**Figure 4A**) prompted us to examine whether pharmacologically targeting mSWI/SNF ATPase activity directly impairs PMN-MDSC motility and function. We found that both AU-24118 and FHD-286 markedly reduced PMN-MDSC motility (**Figure S7E**). Importantly, treatment of mouse PMN-MDSCs with either AU-24118 or FHD-286 markedly abrogated their suppressive activity toward mouse CD8^+^ T cells. This effect was evidenced by preservation of functional CD8_⁺_ T cell frequencies at levels comparable to CD8_⁺_ T cell-only controls, whereas DMSO-treated PMN-MDSCs induced robust effector T cell suppression (**Figure 4H**). A similar phenomenon was observed in human PMN-MDSCs, in which mSWI/SNF antagonism significantly increased the frequency of functional human CD8^+^ T cells relative to suppressive control conditions (**Figure 4I**). Furthermore, in vehicle-treated tumors, S100A9_⁺_ PMN-MDSCs were spatially segregated from CD8_⁺_ T cells, while mSWI/SNF antagonist-treated tumors showed reduced spatial separation between S100A9_⁺_ PMN-MDSCs and CD8_⁺_ T cells (**Figure S7F**). This closer spatial proximity agrees with functional attenuation of PMN-MDSCs following mSWI/SNF antagonism, thereby reducing their exclusionary effect and allowing CD8_⁺_ T cells to localize more readily within PMN-MDSC-containing regions. Collectively, these findings demonstrate that mSWI/SNF antagonism functionally disrupts both the migratory and immunosuppressive programs of PMN-MDSCs.

### mSWI/SNF maintains PU.1/C/EBPβ chromatin programs in suppressive neutrophils

The mSWI/SNF complex is a central regulator that maintains chromatin accessibility [11, 13]. To define the epigenetic mechanisms underlying mSWI/SNF antagonism in PMN-MDSCs, we performed ATAC-seq following treatment with either FHD-286 or AU-24118. As expected, inactivation or degradation of mSWI/SNF ATPases resulted in a global loss of chromatin accessibility across the PMN-MDSC genome (**Figure 5A** and **Figure S8A**). Notably, this response emerged rapidly, within 24 hours, consistent with the concomitant downregulation of PMN-MDSC-associated functional genes (**Figure 4C** and **Figure S7C**). Moreover, AU-24118 and FHD-286 induced highly concordant chromatin remodeling profiles, with approximately 90% overlap in differentially accessible regions (**Figure 5A**), supporting on-target activity and high pharmacologic selectivity of both agents. Motif enrichment analysis of these compacted regions revealed strong enrichment for binding motifs of the master myeloid transcription factors PU.1 and C/EBPβ (**Figure 5B**), which are essential regulators of tumor-driven PMN-MDSC expansion and activation [37, 41, 42]. Consistent with these findings, ChIP-seq analysis demonstrated that mSWI/SNF antagonism markedly diminished the genomic occupancy of both PU.1 and C/EBPβ (**Figure 5C** and **Figure S8C**), with particularly pronounced loss at the PMN-MDSC-associated functional gene loci (**Figure 5D**). Consistently, H3K27 acetylation (H3K27ac), a histone modification marking active promoters and enhancers, was substantially reduced at these loci following mSWI/SNF antagonism (**Figure S8B** and **Figure 5D**). Concordantly, genes exhibiting reduced PU.1 and C/EBPβ promoter occupancy also showed significantly decreased expression by bulk RNA sequencing (**Figure 5E**). Previous studies have reported that mSWI/SNF physically interacts with PU.1 and C/EBPβ to facilitate transcription factor access to chromatin [43-45]. Consistent with this model, reciprocal co-immunoprecipitation revealed physical interactions between the mSWI/SNF ATPase BRG1 and PU.1 and C/EBPβ in PMN_-_MDSCs (**Figure 5F**), supporting a cooperative role for mSWI/SNF and lineage-defining transcription factors in maintaining chromatin accessibility. Together, these data establish that mSWI/SNF sustains the epigenetic and transcriptional programs enforcing PMN-MDSC suppressive identity, and that systemic mSWI/SNF antagonism reprograms PMN-MDSCs by dismantling lineage-defining transcription factor networks and their associated active chromatin landscapes.

**Figure 5.**
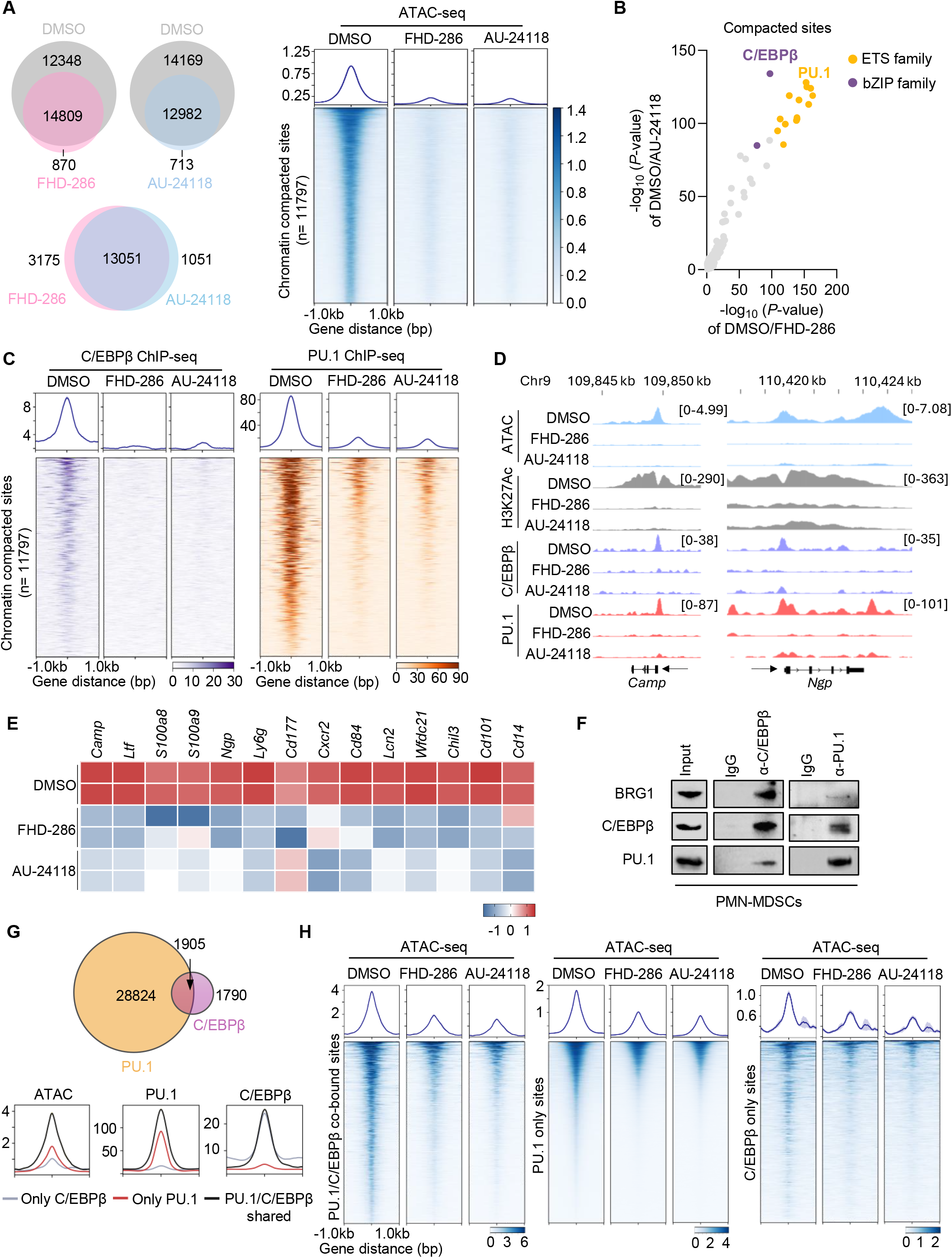
mSWI/SNF antagonism diminishes chromatin accessibility and restricts PU.1 and C/EBPβ binding at PMN-MDSC functional genes. A. Left: Venn diagrams showing overlaps of ATAC-seq peaks between PMN-MDSCs treated for 24 hours with DMSO and FHD-286 (100 nM), DMSO and AU-24118 (3 µM), or FHD-286 and AU-24118. Right: ATAC-seq signal density plots and heatmaps centered on genomic regions exhibiting concordant loss of accessibility following AU-24118 and FHD-286 treatment in PMN-MDSCs, compared with DMSO control. B. Known motif enrichment analysis of compacted chromatin regions following mSWI/SNF antagonism, showing preferential enrichment of ETS family (PU.1) and bZIP family (C/EBPβ) transcription factor motifs (top motifs were highlighted). C. ChIP-seq read-density heatmaps representing C/EBPβ (purple) and PU.1 (orange) at mSWI/SNF antagonists-loss genomic sites in PMN-MDSCs following 24-hour treatment with DMSO, FHD-286 (100 nM), or AU-24118 (3 µM). D. Combined ATAC-seq and ChIP-seq tracks for *Camp* and *Ngp* in PMN-MDSCs with DMSO, FHD-286 (100 nM), or AU-24118 (3 µM) treatments. E. RNA-seq heatmaps for PMN-MDSC signature genes in PMN-MDSCs treated with DMSO, FHD-286 (100 nM), and AU-24118 (3 μM) for 24 hours. *n* = 2 biological replicates. F. Co-immunoprecipitation of BRG1, PU.1, or C/EBPβ in PMN-MDSCs followed by immunoblot for BRG1, PU.1, and C/EBPβ. Data are representative of two independent experiments. G. Top: Venn diagrams showing the overlap of genome-wide PU.1 and C/EBPβ ChIP-seq peaks in PMN-MDSCs. Bottom: Profile plots of ATAC-seq, PU.1 and C/EBPβ ChIP-seq signals at PU.1/ C/EBPβ co-bound, PU.1-only and C/EBPβ-only binding sites in PMN_-_MDSCs. H. ATAC-seq signal density plots and heatmaps at PU.1/ C/EBPβ co-bound sites, PU.1 only sites, and C/EBPβ only sites from PMN-MDSCs treated with DMSO, FHD-286 (100 nM), or AU-24118 (3 μM).

Next, we stratified C/EBPβ and PU.1 binding sites into co-bound regions occupied by both transcription factors and factor-specific regions bound exclusively by either PU.1 or C/EBPβ. Consistent with the role of PU.1 as a master transcription factor, co-bound sites with C/EBPβ have been reported to constitute a much larger fraction of the C/EBPβ cistrome than of the PU.1 cistrome [46]. In PMN-MDSCs, these co-bound sites accounted for approximately 6% of the PU.1 cistrome and 50% of the C/EBPβ cistrome (**Figure 5G**). Importantly, these co-bound regions exhibited substantially higher chromatin accessibility compared with factor-specific sites (**Figure 5G**), indicating cooperative regulation of highly accessible regulatory elements in PMN_-_MDSCs. Notably, mSWI/SNF antagonism led to a marked reduction in chromatin accessibility at PU.1/C/EBPβ co-bound regions, as well as at sites bound exclusively by either transcription factor (**Figure 5H**), indicating that mSWI/SNF activity is broadly required to sustain both cooperative and factor-specific chromatin accessibility programs. Collectively, these data demonstrate that pharmacological mSWI/SNF antagonism collapses PU.1_-_and C/EBPβ_-_dependent chromatin accessibility, thereby extinguishing the transcriptional programs that sustain tumor-driven PMN-MDSC expansion and immunosuppressive function (**Figure 6**).

**Figure 6.**
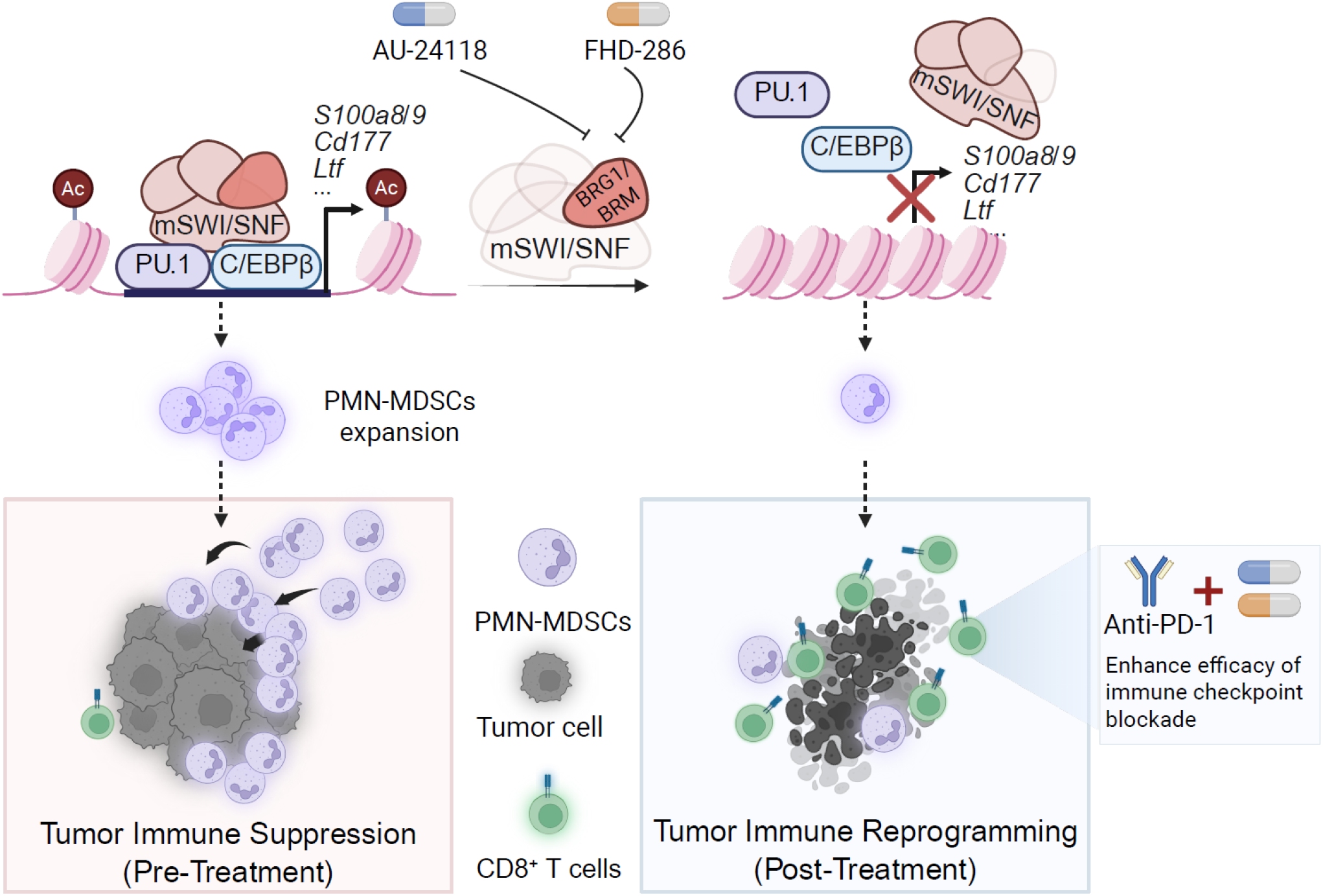
Working model of how mSWI/SNF inhibition suppresses PMN-MDSC expansion and remodels the tumor microenvironment.

## Discussion

This study identifies a druggable chromatin-remodeling vulnerability in tumor-induced suppressive neutrophils. Prior work on mSWI/SNF complexes has largely focused on tumor cell_-_intrinsic transcriptional dependencies or T cell-centric epigenetic programs [13, 15, 47]. By leveraging two mechanistically distinct, systemically bioavailable mSWI/SNF ATPase antagonists together with unbiased immune profiling, genetic perturbation, and immune-cell depletion studies, we identify PMN-MDSCs as the earliest detectable and dominant immune population affected by systemic mSWI/SNF antagonism in tumors. Across multiple syngeneic models, therapeutic activity was lost in immunodeficient hosts yet retained against tumors lacking *Smarca2/4*, indicating that tumor control was not driven primarily by tumor-cell-intrinsic mSWI/SNF dependence. Instead, mSWI/SNF antagonism rapidly reduced suppressive PMN_-_MDSCs, followed by expansion and functional activation of CD8^+^ T cells. Anti-Ly6G depletion, CD8^+^ T cell depletion, and *Smarca4*^f/f^*S100a8*^Cre^ genetic studies further support a model in which mSWI/SNF antagonism acts through a PMN-MDSC-CD8^+^ T cell immune axis. These findings establish mSWI/SNF-dependent chromatin remodeling as a previously unrecognized regulator of tumor-associated suppressive neutrophils.

PMN-MDSCs represent a dominant mechanism of immune evasion across many solid tumors [4, 48, 49]. These pathologically activated neutrophil-like cells arise through tumor-induced emergency granulopoiesis and impair antitumor immunity by suppressing effector lymphocyte function [49]. PMN-MDSCs are enriched in immune checkpoint blockade-refractory tumors, where their accumulation correlates with poor prognosis and inferior responses to immunotherapy [7, 50, 51]. Although substantial effort has been devoted to targeting PMN_-_MDSCs, primarily through inhibition of chemokine-mediated trafficking, survival pathways, or suppressive effector mechanisms, these approaches have often produced incomplete or transient activity [9, 52]. Even advanced clinical efforts, including tasquinimod, failed to improve overall survival in a phase III setting [8]. In direct comparison in the tested model, mSWI/SNF antagonism reduced PMN-MDSC abundance and controlled tumor growth more effectively than tasquinimod, supporting chromatin remodeling as a proximal point of intervention in PMN_-_MDSC-driven immune suppression.

Mechanistically, our work identifies a state-specific epigenetic dependency that sustains the pathological identity of PMN-MDSCs. Integrated analyses of chromatin accessibility, transcription factor occupancy, histone acetylation, and gene expression show that tumor_-_associated PMN-MDSCs require continuous mSWI/SNF ATPase activity to maintain PU.1- and C/EBPβ-associated regulatory programs. Although PU.1 is a master regulator of myeloid lineage commitment, prior studies have shown that PU.1-bound regulatory elements differ in their dependence on chromatin remodeling complexes, with dynamically activated enhancers exhibiting heightened mSWI/SNF sensitivity [53, 54]. Our data extend these observations by showing that tumor-induced PMN-MDSCs are particularly dependent on mSWI/SNF-mediated accessibility at PU.1- and C/EBPβ-occupied regulatory regions. Pharmacologic mSWI/SNF antagonism caused a rapid and concordant collapse of chromatin accessibility, H3K27 acetylation, and PU.1/C/EBPβ occupancy at regulatory elements linked to PMN-MDSC expansion, migration, and suppressive function. Thus, mSWI/SNF antagonism does not merely attenuate individual PMN-MDSC effector pathways; it dismantles the chromatin architecture that maintains the suppressive-neutrophil state.

The state-specific nature of this vulnerability may help explain the therapeutic window observed in these models. Over the treatment intervals examined, systemic mSWI/SNF antagonism preferentially depleted tumor-associated PMN-MDSCs while sparing other major immune populations and preserving CD8^+^ T cell activation. In human systems, mSWI/SNF antagonism blocked tumor explant-induced expansion of LOX-1^+^ PMN-MDSCs while sparing LOX-1^-^ neutrophils, suggesting that mSWI/SNF sensitivity is linked to pathological neutrophil activation rather than neutrophil identity alone. These findings are particularly important because genetic disruption of mSWI/SNF components can impair lymphocyte development, whereas transient, dose-controlled pharmacologic antagonism in our models preserved and restored effector immune function. Together with the absence of significant treatment-associated weight loss in our studies and prior preclinical work with AU-24118 and FHD-286 [15, 19, 20], these data support a therapeutic window for systemic mSWI/SNF antagonism in PMN-MDSC-rich tumors.

These findings have several translational implications. First, they suggest that PMN-MDSC-rich tumors, particularly those refractory to checkpoint blockade, may be especially susceptible to mSWI/SNF antagonism. Second, they provide a rationale for combining mSWI/SNF ATPase antagonists with PD-1 blockade, because depletion of suppressive neutrophils removes a barrier to CD8^+^ T cell expansion and function. Third, they nominate potential pharmacodynamic biomarkers, including LOX-1^+^ PMN-MDSCs in human samples, S100A8/S100A9 and CD177 expression, PU.1/C/EBPβ-associated chromatin accessibility, and spatial redistribution of PMN_-_MDSCs and CD8^+^ T cells. The clinical advancement of mSWI/SNF ATPase antagonists further supports evaluation of this concept in human cancers [55], although dosing schedules, combination regimens, and immune-monitoring strategies will need to be optimized.

Together, these findings establish suppressive-neutrophil chromatin remodeling as a therapeutic vulnerability in cancer. Tumor-induced PMN-MDSCs are not merely inflammatory byproducts of malignancy, but pathological immune states maintained by a druggable mSWI/SNF-dependent chromatin program. Dismantling this dependency restores CD8_⁺_ T cell immunity, enhances checkpoint blockade, and provides a framework for chromatin-based immunotherapy in PMN_-_MDSC-rich, checkpoint-refractory tumors.

## Methods

### Cell lines and compounds

B16-F10, CT26, A20, 4T1 were originally obtained from American Type Culture Collection (ATCC). MC38 was provided by Dr. Walter Storkus. All cells were genotyped to confirm their identity at the Labcorp Cell Line Testing division (Burlington) for authentication and tested routinely for Mycoplasma contamination. Gibco ATCC modified RPMI 1640 + 10% FBS (Thermo Fisher Scientific) were used for B16-F10, CT26, A20 and 4T1. Gibco DMEM + 10% FBS (Thermo Fisher Scientific) were used for MC38. All cells were cultured in a humidified incubator at 37 °C and 5% CO_2_. AU-24118 and AU-15330 were provided by Aurigene. FHD-286 was provided by Foghorn Therapeutics. Tasquinimod was purchased from MedChem Express.

### Cell proliferation

Cells were plated directly into 96-well plates and were placed into an IncuCyte Live Cell Analysis Imaging System S3 (Sartorius) in a humidified incubator at 37 °C and 5% CO_2_. Phase object confluence (percentage area) was acquired every 4 h using the 10x objective.

### qRT-PCR

Total RNA was extracted from cells using QIAzol Lysis Reagent (QIAGEN) according to the manufacturer’s instructions. The RNA was then eluted in RNase-free water, and its concentration and purity were assessed using a NanoDrop spectrophotometer.

For quantitative PCR (qPCR) analysis, cDNA was synthesized using the extracted RNA as a template by using LunaScript RT SuperMix Kit (NEB, E3010L). The resulting cDNA was then used for qPCR using Fast SYBR Green Master Mix (Applied biosystems, 4385612). The qPCR reactions were performed in a 384-well format on QuantStudio 5 or 7 Pro system (Thermo Fisher Scientific), and the data were analyzed using the 2^−ΔΔCT^ method to quantify gene expression levels, normalizing to the expression of the *Actb* gene. Information for the primers used in this study is listed in **Supplementary Table 2**.

### CRISPR–Cas9-mediated gene knockout

Guide RNAs (sgRNAs) targeting the early exons of mouse *Smarca4/Brg1* or *Smarca2/Brm* were checked for off-target prediction using Off-Spotter (https://cm.jefferson.edu/Off-Spotter/). Those sgRNAs with weak off-target potential together with non-targeting sgRNA were cloned into lentiCRISPR v2 plasmid (Addgene; #52961) according to published literature. B16-F10 or MC38 cells were transiently transfected with lentiCRISPR v2 encoding non-targeting or pool of *Smarca4/Brg1* and *Smarca2/Brm* targeting sgRNAs. After 24 hours transfection, cells were selected with puromycin, and then single cells were plated into 96-well dishes with a cell sorter (Sony SH800S). Sanger sequencing and Western blot were performed to examine the depletion of target genes. The sgRNA sequences are listed in **Supplementary Table 3**.

### Animal experiments

All mouse studies conducted in this study were approved by the U-M Institutional Animal Care and Use Committee. *Smarca4^f/f^*mice (B6;129S2-*Smarca4^tm1Pcn^*/Mmnc) were obtained from the Mutant Mouse Resource and Research Centers (MMRRC) repository (MMRRC ID: 36548). *S100a8*-Cre (Stock# 021614) mice were obtained from the Jackson Laboratory (Bar Harbor, ME). Animals were backcrossed, interbred, and maintained on a C57BL/6J background. Female C57BL/6J mice aged 6-8 weeks were purchased from The Jackson Laboratory (Stock# 000664) and used for MC38 and B16-F10 models. Female BALB/c mice aged 6-8 weeks were purchased from Charles River and used for CT26, 4T1, and A20 models. Female NOD.Cg-*Prkdc^scid^ Il2rg^tm1Wjl^*/SzJ (NSG) mice aged 6-8 weeks were purchased from the Jackson Laboratory (Stock#005557) and used for all the models. Subcutaneous tumor models were established by inoculating 3 × 10^6^ MC38 cells, 5 × 10^5^ CT26 cells, 3 × 10^5^ B16-F10 cells, or 3 × 10^6^ A20 cells at both flanks of their syngeneic immunocompetent mice or NSG mice. The orthotopic mammary fatpad breast tumor model was established by inoculating 3 × 10^5^ 4T1 cells in the mammary fat pad of BALB/c and NSG mice. Tumor volume was measured 6 to 8 days after tumor cell inoculation every 2 to 4 days, using calipers. Tumor volume was calculated using the formula V = (W^2^× L)/2, where W and L represent the minor and major tumor axes, respectively. Animals were pathogen free and were housed on a 12-h light -12-h dark cycle. Mice were euthanized when total tumor burden reached 2 cm in diameter or earlier if humane end point criteria were observed.

### Drug formulation and in vivo treatments

AU-24118 was added in one part of PEG200 and then sonicated and vortexed until completely dissolved. Five parts of 10% D-α-Tocopherol polyethylene glycol 1000 succinate were next added, followed by adding four parts of 1% Tween-80. The solution was vortexed until homogeneous. FHD-286 was dissolved in 20% (2-Hydroxypropyl)-beta-cyclodextrin and sonicated until homogeneous. Tasquinimod was first dissolved in DMSO, then mixed with 40% PEG300, 5% Tween-80, and 45% water.

Mice were randomized when tumors reached 30 to 100 mm^3^. AU-24118 was administered orally at 7.5 mg/kg once daily, FHD-286 at 1.5 mg/kg once daily, and tasquinimod at 30 mg/kg once daily. Anti-PD-1 (clone RMP1-14, Bio X Cell) and its isotype control were administered intraperitoneally at 100 μg per mouse for MC38 tumor models and at 200 μg per mouse for other models two times a week. For depletion of PMN-MDSCs, anti-mouse Ly6G (clone 1A8, Bio X Cell) or the corresponding isotype controls were administrated intraperitoneally 1 day before treatment or co-treatment of vehicle, AU-24118, or FHD-286, at 200 μg per mouse every other day. For depletion of T cells, anti-mouse CD8α (clone 2.43, Bio X Cell) or the corresponding isotype controls were administrated intraperitoneally 1 day before tumor cell inoculation or co-treatment with vehicle, AU-24118, or FHD-286, at 400 μg per mouse, and subsequently 100 μg per mouse one time every 3 days.

### Flow cytometry analysis

Single-cell suspensions were prepared from fresh mouse tumor tissues or spleens as described previously [56]. Cells were stained with Zombie Green (BioLegend, 423112) in PBS, blocked with anti-mouse CD16/32 (BioLegend, 156604) in MACS (PBS containing 2% FBS and 2 mM EDTA), and subsequently stained with surface antibodies in MACS for 12 minutes at room temperature. To assess intracellular cytokine production in T cells, cells were cultured for 4 h in the presence of phorbol 12-myristate 13-acetate (200 ng/mL; Sigma-Aldrich), ionomycin (1000 ng/mL; Sigma-Aldrich), 27.5 µM β-mercaptoethanol (Gibco), 1× monensin (Thermo fisher scientific) and 1×brefeldin A (BD Biosciences). Cells were fixed and permeabilized by the Foxp3/Transcription Factor Staining Buffer Set (Thermo Fisher Scientific, 00-5523-00). Cellular phenotypes were assessed and analyzed on a flow cytometer (BD LSRFortessa cell analyzer) with Absolute Counting Beads (Thermo Fisher Scientific, C36950) added for quantification. Antibodies against the following mouse antigens were used: anti-CD45 (Biolegend, 103132); anti-CD3 (BioLegend, 100216); anti-CD19 (Biolegend, 115508); anti-NK1.1 (Biolegend, 108714); anti-CD90.2 (BD Biosciences, 564365); anti-CD8 (BioLegend, 100742); anti-CD4 (BD Biosciences, 563106); anti-CD11b (Biolegend, 101208); anti-F4/80 (BD Biosciences, 565613); anti-CD11c (Thermo Fisher Scientific, 25-0114-82); anti-I-A/I-E (BD Biosciences, 562564); anti_-_Ly-6G/Ly-6C (Gr-1) (Biolegend, 108422); anti-Ly6C (Biolegend, 128033); anti-Ly6G (Biolegend, 127624); anti-Ki67 (Thermo Fisher Scientific, 56-5698-82); anti-granzyme B (BioLegend, 372208); anti-IFN-γ (BD Biosciences, 562303); anti-TNF-α (Biolegend, 502930). The strategy for immune cell gating is presented in **Supplementary Figure 2B**.

### scRNA-seq and data analysis

Fresh mouse tumors were processed into single-cell suspensions as described previously [56]. Cells were first stained with Zombie Green viability dye (BioLegend, 423112) in PBS, followed by Fc receptor blocking with anti-mouse CD16/32 antibody (BioLegend, 156604) in MACS buffer (PBS containing 2% FBS and 2 mM EDTA). Cells were then stained with PerCP/Cy5.5_-_conjugated anti-mouse CD45 antibody (BioLegend, 103132) in MACS buffer for 12 min at room temperature. Viable CD45^+^ cells were purified using a Sony SH800S cell sorter. Single-cell RNA-seq libraries were generated using the Chromium Next GEM Single Cell 3 ’ HT Kit v3.1 Dual Index (10x Genomics) according to the manufacturer’s protocol and sequenced on an Illumina NovaSeq platform. After data processing and quality control, the final integrated dataset comprised 31,588 CD45^+^ single-cell transcriptomes across all experimental conditions. Per-sample FASTQ files were generated from the raw base call files using 10x Cell Ranger v.9.0.0 *mkfastq* [57]. Read demultiplexing, alignment, and gene expression quantification were conducted using the 10X Genomics Cell Ranger count pipeline v.9.0.0 with the pre-built mouse reference genome mm10. Counts were adjusted for background contamination using the R package SoupX v1.6.2 [58]. Cells were excluded from downstream analyses if they were identified as putative doublets by scDblFinder, exhibited low gene complexity with fewer than 200 detected genes, contained a mitochondrial transcript fraction exceeding 20%, or had fewer than 500 total transcript counts. The samples were processed using the R package Seurat (v5) standard workflow methods as described [59]. Cluster markers were identified using *FindAllMarkers*, visualized with DotPlot, and used to subcluster the data into lymphoid and myeloid subclusters [60]. The subclusters were further processed using Seurat, and cell clusters were manually annotated using marker genes of the subclustered results. T cell clusters were subsetted and subjected to independent reanalysis to enable higher-resolution identification of transcriptionally distinct subpopulations. The resulting subclusters were annotated based on their top 10 differentially expressed genes, complemented by manual assessment of selected canonical markers defining T cell lineage and functional states.

### Xenium In Situ (10x Genomics) workflow

Mouse tissue microarrays (TMAs) generated from formalin-fixed, paraffin-embedded (FFPE) syngeneic mouse tumor models were sectioned at 5 µm thickness and mounted onto Xenium slides. Slides were baked at 60 °C for 2 hours, followed by deparaffinization and tissue decrosslinking to expose target RNA, according to standard 10x Genomics protocols (Protocol CG000580). Tissues were hybridized with the Xenium Prime 5K Mouse Pan Tissue & Pathways Panel (PN-1000725) and a 100-gene mouse-specific custom add-on panel (Xenium Prime 5K Custom Add-On Panel, 51–100 genes, PN-1000766, Design ID# X6JM38) (**Supplementary Table 1**). Rolling circle amplification (RCA) was performed to enzymatically generate hundreds of copies of the gene-specific barcode per RNA target in situ. Cell segmentation and autofluorescence quenching were performed according to manufacturer’s instructions (Protocol CG000760), followed by DAPI nuclear staining. Multiplexed in situ RNA profiling was performed using the 10x Genomics Xenium Analyzer (Protocol CG000584 RevK).

Outputs were reprocessed using Xenium Ranger to restrict cell segmentation to nuclei staining using an expansion distance of 1 µm and a DAPI intensity filter of 400. Xenium Ranger outputs were joined processed using Seurat v5.2.1 and R v4.4. Each individual TMA core was assessed for tumor viability and the cells outside of viable regions were excluded from downstream analyses. Cells with less than 50 detected transcripts were also excluded from downstream analyses. The remaining cells were processed using a standard Seurat pipeline, with a FindClusters resolution of 0.3. Broad cell type annotation was performed using the top ten markers for each cluster from FindAllMarkers. Myeloid and lymphocyte clusters were separated and reprocessed using a standard pipeline with a lowered number of dimensions (dims = 15). Functional annotation of these subclusters was done using RCTD [61] with our previously described single-cell RNA-seq data as a reference. Final annotation was based on gene markers from the subset analysis and results from RCTD. PMN-MDSC related transcript densities were visualized using Xenium Explorer 4.

### Immunofluorescence

Immunofluorescence (IF) was performed for BRG1 (Abcam, ab108318) and BRM (Millipore Sigma, HPA029981) on 5-μm-thick formalin-fixed, paraffin-embedded (FFPE) tissue sections using the Ventana ULTRA automated slide staining system (Roche-Ventana Medical Systems) with the anti-rabbit OmniView HRP (Roche-Ventana, 760-4311) and Cy5 Tyramide/Amplification reagent (catalog no. 760-238, Roche-Ventana) to develop a fluorescent signature, followed by DAPI (Prolong gold anti-fade, P36931, Invitrogen/Fisher-Scientific) for counterstaining nuclei, as previously described [62]. IF staining was evaluated using an EVOS 7000 system (Fisher Scientific). Staining was independently assessed by two study pathologists (R.P. and R.M) at ×100 and ×200 magnification to assess and quantify biomarker expression.

Multiplex immunofluorescence was performed using the fully automated COMET™ platform (Lunaphore Technologies, Tolochenaz, Switzerland). The antibody panel included anti-S100A9 (Cell Signaling Technology, 73425S), anti-CD8 (Cell Signaling Technology, 98941S), anti-BRG1 (Abcam, ab108318), anti-BRM (Millipore Sigma, HPA029981), and anti-CD177 (Cell Signaling Technology, 28740S). Raw images were visualized and exported using HORIZON^TM^ viewer software. The raw data were comprised of single-cell protein expression values across multiple markers along with associated spatial coordinates. Markers were split into biologically relevant panels for downstream visualization and analysis. Outliers were removed separately for each marker within each sample by excluding cells with expression values outside ±3 median absolute deviations (MAD) from the sample-specific median. This approach reduces the influence of technical or biological outliers while preserving most observations for downstream statistical analysis. Following outlier removal, expression values were log1p-transformed to stabilize variance and reduce the influence of skewed distributions. The nearest neighbor analysis was performed in each sample using RANN package (version 2.6.2) in R (version 4.5.3). This analysis quantifies the distance from each cell in a chosen reference population to its closest cell in a target population.

### Immunoblotting

Cells were washed once in PBS and scraped into RIPA lysis buffer containing protease and phosphatase inhibitors (78440, Thermo Fisher Scientific). After sonication and removal of debris by centrifugation, protein concentration was determined via Rapid Gold BCA Protein Assay (Thermo Fisher Scientific, A55860) and samples were mixed with LDS sample buffer (Thermo Fisher Scientific, NP0007) with reducing reagents (Thermo Fisher Scientific, NP0009), followed by a 10 min incubation at 70 °C. Proteins were resolved by either NuPAGE 3-8% Tris-Acetate or 4-12% Bis-Tris Protein Gels (Thermo Fisher Scientific), followed by transfer to 0.45 μm PVDF membranes (Millipore). Membranes were blocked in 5% non-fat milk in TBST and probed overnight at 4 °C with the appropriate antibody. Membranes were then incubated with horseradish peroxidase-linked secondary antibody and chemiluminescent signal was detected with an Odyssey CLx Imager (LI-COR Biosciences) or ChemiDoc XRS+ Imaging System (Bio_-_Rad). The following antibodies were used: BRG1 (1:1000; Cell Signaling Technology, 52251); BRM (1:1000; Abcam, ab240648); PBRM1 (1:1000; Bethyl Laboratories, A301-591A-A); vinculin (1:1000; Cell Signaling Technology, 18799S); C/EBPβ (1:1000; Abcam, ab32358); PU.1 (1:1000; Cell Signaling Technology, 2258S); ECL peroxidase (HRP) anti-rabbit IgG (1:4000; Cytiva, NA934-1ML); ECL peroxidase (HRP) anti-mouse IgG (1:2000; Cytiva, NA931-1ML).

### PMN-MDSCs isolation

For mouse PMN-MDSC isolation, spleens or tumors were harvested and dissociated to generate single-cell suspensions. Red blood cells (RBC) were removed using RBC lysis buffer (BioLegend, 420301). PMN-MDSCs were then isolated using the Myeloid-Derived Suppressor Cell Isolation Kit (Miltenyi Biotec, 130-094-538) according to the manufacturer’s recommendations, or by cell sorting (live CD45^+^ CD11b^+^ Ly6G^+^).

For human PMN-MDSC isolation, peripheral blood samples were obtained from patients with prostate cancers or breast cancers from the MiOncoSeq Clinical Laboratory at The Michigan Center for Translational Pathology (MCTP) under the approval of the University of Michigan Institutional Review Board. Written informed consent was obtained from all participants prior to sample collection. Samples were de-identified before processing and analysis. Blood was added to the Ficoll, and PMBCs layers were collected. Red blood cells were removed using RBC lysis buffer. PMN-MDSCs were sorted by a cell sorter (live CD66b^+^ LOX-1^+^).

### Co-immunoprecipitation

PMN-MDSCs lysates were prepared in Pierce IP lysis buffer (Thermo Fisher Scientific, 87788) supplemented with protease and phosphatase inhibitors (Thermo Fisher Scientific, 78440). Cell lysates were sonicated and centrifuged for 10 mins at maximum speed. The supernatant was precleared by Dynabeads Protein G (Thermo Fisher Scientific, 10004D) with the control IgG for 2 hours at 4°C. 1% input sample was kept. Lysates were incubated with BRG1 (Abcam, ab110641), PU.1 (Invitrogen, PA5-17505), or C/EBPβ (Proteintech, 23431-1-AP) antibody overnight at 4°C. The following day, Dynabeads Protein G were added and incubated for an additional 2 h at 4°C. Beads were subsequently washed four times with IP lysis buffer, and bound proteins were eluted for immunoblot analysis.

### TES preparation

Tumor explant supernatant (TES) was generated following an established protocol [59]. Briefly, approximately 5 g of freshly excised 4T1, B16-F10, or human tumor tissues were finely chopped and cultured in 20 mL of complete ATCC modified RPMI-1640 medium. After 24 hours incubation, conditioned medium was harvested, filtered by passing through a 0.22-µm filter, snap-frozen, and stored at -80 °C until use. Human tumors were collected from the Cooperative Human Tissue Network with patient consent. The use of the tissue samples was approved by the Institutional Review Board at the University of Michigan.

### T cell suppression assay

Isolated mouse or human PMN-MDSCs were pre-treated with DMSO, FHD-286 (100 nM), AU_-_24118 (3 µM), or tasquinimod (30 µM) for 24 hours. Mouse or human CD8^+^ T cells were isolated from mouse spleens or frozen human PBMC (Lonza) by the EasySep™ Mouse/human CD8^+^ T Cell Isolation Kit (Stemcell), followed by activation for 2 days using the T Cell Activation/Expansion Kit (Miltenyi Biotec, 130-093-627) for mouse CD8^+^ T Cell or T cell TRANS Act (Miltenyi Biotec, 130-128-758) for human CD8^+^ T Cell. PMN-MDSCs were then co-cultured with CD8^+^ T cells for 72 hours. IFN-γ was then measured on CD8^+^ T cells by flow cytometry.

### Migration assay

Cell migration was assessed using a Transwell assay with 5-µm pore inserts (Corning). PMN-MDSCs were isolated and pre-exposed for 24 h to DMSO, FHD-286 (100 nM), AU-24118 (3 µM), or AU-15330 (3 µM) prior to being seeded into the upper chamber. RPMI-1640 medium was added to the lower chamber. Cells were allowed to migrate for 2 h, after which cells in the lower chamber were collected and quantified using an automated cell counter. Migration efficiency was calculated as the ratio of migrated cells to the total number of cells initially seeded.

### Bulk RNA-seq and data analysis

RNA-seq libraries were prepared using 200-1,000 ng of total RNA by using KAPA RNA HyperPrep Kit with RiboErase (Roche Sequencing Solutions, 08098140702) following the user’s manual. Libraries were prepared via enzymatic rRNA depletion, heat fragmentation (200–300 bp), and cDNA synthesis. Resulting double-stranded cDNAs were ligated with NEB adapters and amplified using KAPA HiFi HotStart mix and dual barcodes. Library quality was assessed with the Agilent Bioanalyzer using DNA 1000 Kit (Agilent Technologies; catalog #5067-1504) and then sequenced by NovaSeq 6000.

Libraries passing quality control were trimmed of sequencing adapters and aligned onto the GRCm38/mm10 mouse reference genome. Data analysis was performed with packages limma [63, 64] and edgeR [65]. Gene set enrichment analysis was performed as described [56]. Additional plotting and Gene Ontology enrichment analysis was performed using clusterprofiler [66].

### ATAC-seq and data analysis

ATAC-seq was performed as previously described [15]. In brief, 0.5 × 10^6^ PMN-MDSCs treated with AU-24118 and FHD-286 were washed in cold PBS once and resuspended in RSB buffer supplemented with NP-40, Tween-20, protease inhibitor, and digitonin. This single-cell suspension was incubated on ice for 5 min. The lysing process was quenched by adding double the same volume of RSB buffer with Tween-20 and protease inhibitor. The lysate was centrifuged at 500g for 5 min at 4°C to remove the supernatant, followed by incubating with 2 μl Tn5 enzyme for 30 min at 37°C (20034198, Illumina Tagment DNA Enzyme and Buffer Kit). Samples were immediately purified by Qiagen minElute column and PCR-amplified with the Next High-Fidelity 2X PCR Master Mix (NEB, M0541L) following the original protocol. Optimal PCR cycles were determined via qPCR to avoid over-amplification. The amplified library was further purified by Qiagen minElute column and Ampure beads. ATAC-seq libraries were sequenced on the Illumina HiSeq 2500 or NovaSeq using a 2 × 50 nucleotide paired-end read length with sequence depth of 30-35 million reads per sample.

Sequencing of ATAC-seq libraries generated fastq files, which were processed as described previously [15, 67]. To summarize, these fastqs were initially processed using Trimmomatic (v.0.39)[68] for trimming. These files were then aligned to the GRCm38/mm10 mouse genome using bwa mem (v.0.7.17-r1198-dirty)[69], and the alignments were converted to binary format with SAMtools (v.1.9) [70]. We next eliminated reads from mitochondrial DNA and duplicated reads, using SAMtools and Picard MarkDuplicates (v.2.9) [71]. Peaks in the ATAC-seq data were identified using MACS2 (v.2.1.1.20160309) [72]. Finally, conversion of data to bigwig format was accomplished using the UCSC tool wigtoBigwig [73]. Known motif enrichment analyses were performed using the HOMER (version v4.11.1) [46] suite of algorithms. Comparisons between samples to determine the sites present in DMSO but lost upon AU-24118 or FHD-286 treatment were conducted using the R packages ChIPseeker [74] and ChIPpeakAnno (version 3.0.0) [75, 76]. These reduced accessibility sites were then plotted as read density heatmaps using deepTools [77].

### ChIP-seq and data analysis

ChIP experiments were carried out using the ideal ChIP-seq kit for TFs or histones (Diagenode) as per the manufacturer’s protocol. In brief, PMN-MDSCs (2 × 10^6^ cells for H3K27Ac and PU.1 ChIP-seq, 6 × 10^6^ cells for C/EBPβ ChIP-seq) were washed with 1× PBS, followed by cross-linking for 10 min in 1% formaldehyde solution. Crosslinking was terminated by the addition of 1/10 volume 1.25 M glycine for 5 min at room temperature followed by cell lysis and sonication, resulting in an average chromatin fragment size of 200-600 bp. Fragmented chromatin was used for each ChIP reaction with individual antibody (8μg for transcription factor or 1 μg for histone) with overnight incubation at 4°C. ChIP DNA was de-crosslinked and purified according to the standard protocol. Purified 1-20 ng DNA was then prepared for sequencing as described previously [15]. Libraries were quantified and quality checked using the Bioanalyzer 2100 (Agilent) and sequenced on the Illumina HiSeq 2500 or NovaSeq Sequencer (2 × 150-nucleotide read length with sequence depth of 25-35 M paired reads).

ChIP-seq data were analyzed as previously described [15, 67]. In short, ChIP-seq data analysis started with trimming, and reads were aligned to GRCm38/mm10 mouse genome reference. Alignments were filtered using SAMtools [70] and Picard MarkDuplicates [71]. Peaks were called with MACS2 [72] and compared between datasets using ChipSeeker and ChipPeakAnno [74-76]. Finally, ChIP peak profile plots and read-density heatmaps were generated using deepTools [77].

### Statistics

All data points were derived from distinct samples. Graphpad Prism 10 was used to generate graphs and calculate statistics using appropriate statistical tests depending on the data. Means ± SD or ± SEM were used for data presentation, and details of the statistical analysis are provided in the corresponding figure legends. A *P* value less than 0.05 was used to indicate statistical significance. All statistics were adjusted with Bonferroni correction. For GSEA, statistical significance was assessed by permutation-based testing, and the normalized enrichment score (NES) and adjusted *P* value are shown.

## Author Contributions

F.Y., Y.B., L.X., and A.M.C. conceived and designed the studies. F.Y., R.X., and M.G. performed all the in vitro experiments with assistance from W.Z., Y.C., L.V., I.K., J.L., Y.L., and A.C.; F.Y. and Y.B. performed all the in vivo experiments with assistance from Y.Q., J.C.T., W.Z., and M.K.; R.M., R.P., S.M., R.S., and A.R. performed all the histology-based experiments and analysis; X.C. generated next-generation sequencing libraries and performed the sequencing; G.C., E.Y., and Y.Z. carried out all bioinformatics analyses; S.S., C.A., and M.R. contributed to the discovery of AU-24118 and AU-15330 compounds; J.C.P. and M.H. contributed to the discovery of FHD-286 compound; F.Y., Y.B., S.J.M., and A.M.C. wrote the manuscript and organized the final figures.

## Acknowledgements

We thank Xiaoju Wang, Sumit Das, Jiayi Zhou, Rui Wang, Shannon Van Aken, Amanda Miller, Christine Caldwell-Smith, Ashley Wolfe, Mingyu Park, Sophie Yeung, Yang Zheng and Parker Schanen from the Michigan Center for Translational Pathology at the University of Michigan. This work was supported by the following: National Cancer Institute Specialized Programs of Research Excellence grant (P50CA186786, A.M.C.), Trailsend Foundation (A.M.C., L.X., Y.Q.), Prostate Cancer Foundation Challenge Award (A.M.C.), Myeloma Solutions Fund (A.M.C., L.X.). A.M.C. is a Howard Hughes Medical Institute Investigator, A. Alfred Taubman Scholar, and American Cancer Society Professor.

## Supplemental Figure Legends

**Figure S1. Antitumor efficacy and tolerability of mSWI/SNF antagonists in syngeneic tumor models.** A. Representative immunofluorescence (IF) images (left) and quantification (right) assessing BRG1 and BRM levels in A20 subcutaneous (s.c.) tumors from NSG mice. Scale bar: 50 μm. Data were statistically assessed using two-tailed Student’s *t* test.

B. Survival analysis of MC38 s.c. tumors in mice treated with vehicle, AU-24118, or FHD_-_286 in combination with IgG or anti-PD-1 (*n* = 6–7 mice per group).

C. Body weight of syngeneic mice bearing A20 (left), MC38 (middle), or CT26 (right) s.c. tumors and treated with the indicated agents (*n* = 6–8 mice per group).

All data are presented as mean ± SEM and are representative of two independent experiments.

**Figure S2. Immune profiling of mSWI/SNF antagonist-treated tumors by scRNA-seq and flow cytometry.** A. Left: Heatmap showing top differentially expressed genes in each of the indicated clusters among CD45^+^ leukocytes from single cell RNA-sequencing. Right: Heatmap showing top differentially expressed genes in each of the indicated subclusters among CD8^+^ T cells. Three representative genes for each cluster are shown next to the heatmaps. B. Gating strategy for live lymphocytes, intracellular markers in CD8^+^ T cells, or myeloid cells.

**Figure S3. mSWI/SNF antagonists reprogram the immune suppressive tumor microenvironment in various syngeneic models.** A. Flow cytometric analysis of absolute numbers of the indicated immune cells in MC38 subcutaneous (s.c.) tumors from C57BL/6 treated with vehicle, AU-24118, and FHD-286 for 10 days (*n* = 7 mice per group). Representative contour plots showing the proportional change of TNF-α^+^ CD8^+^ T cells are shown on the left. B-E. Flow cytometric analysis of absolute numbers of the indicated immune cells in the indicated models treated with vehicle (gray), AU-24118 (blue), and FHD-286 (pink) for ten days (*n* = 5–7 mice per group). Data are presented as box and whisker plots and were statistically assessed using two-way analysis of variance. All data are representative of two independent experiments.

**Figure S4. Immune profiling of mSWI/SNF antagonist-treated tumors by Xenium In Situ and multiplex immunofluorescence.** A. Dot plot showing the top differentially expressed genes for annotating cell clusters in Xenium In Situ on 4T1 tumors. B. Dot plot showing the expression of representative genes and functional markers used for annotating CD8^+^ T cell subclusters in A. C. UMAP visualization of the clustering results in A and B.

**Figure S5. mSWI/SNF antagonism inhibits tumor progression in a PMN-MDSC-dependent manner.** A. Tumor volume changes over time of CT26 subcutaneous (s.c.) tumors in BALB/c treated with vehicle, AU-24118, or FHD-286 in combination with IgG or anti-Ly6G (*n* =7–10 mice per group). α-Ly6G: anti-Ly6G. B. Tumor volume changes over time of B16-F10 s.c. tumors in C57BL/6 (left, *n* = 6–7 mice per group), or CT26 s.c. tumors in BALB/c (right, *n* = 8–10 mice per group) treated with vehicle, AU-24118, or FHD-286 in combination with IgG or anti-CD8. α-CD8: anti-CD8. C. Flow cytometric analysis of absolute numbers of PMN-MDSCs and CD8^+^ T cells on B16_-_F10 s.c. tumors derived from *Smarca4^f/f^* or *Smarca4^f/f^S100a8^cre^*mice (*n* = 4 mice per group). D. Left: Tumor volume changes over time of B16-F10 s.c. tumors in C57BL/6 treated with vehicle, tasquinimod, AU-24118, or FHD-286 (*n* = 8 mice per group). Right: Flow cytometric analysis of absolute numbers of PMN-MDSCs and IFNγ^+^ CD8^+^ T cells in B16_-_F10 s.c. tumors following treatment with vehicle, AU-24118, or FHD-286 for eight days (*n* = 5 mice per group). Data are presented as mean ± SEM in A, B, and D (left), or as box and whisker plots in C and D (middle and right). All data were statistically assessed using two-way analysis of variance and are representative of two independent experiments. Bonferroni’s correction is applied for multiple comparisons in A, B and D.

**Figure S6. mSWI/SNF antagonism downregulates PMN-MDSC-associated genes.** A. Gene Ontology pathways for neutrophil extravasation enriched in bulk RNA-sequencing profiles of tumor-infiltrating PMN-MDSCs from mice treated with FHD-286 or AU-24118 compared with vehicle (Veh.). PMN-MDSCs were isolated from five mice per group. B. Representative COMET™ multiplex immunofluorescence images showing S100A9 and BRG1 or S100A9 and BRM co-expression in 4T1 mammary fat pad tumors following treatment with vehicle, AU-24118, or FHD-286. Scale bar, 10 µm. Right: Quantification shows the single-cell protein expression of BRG1 and BRM in S100A9^+^ cells across treatment groups. C. Representative COMET™ multiplex immunofluorescence images (left) and quantification (right) of COMET™ multiplex immunofluorescence showing S100A9 and CD177 expression in S100A9^+^ PMN-MDSCs within 4T1 mammary fat pad tumors following treatment with vehicle, AU-24118, or FHD-286. Scale bar, 10 µm. Data are presented as box and whisker plots in B and C, and were statistically assessed using two-way analysis of variance.

**Figure S7. mSWI/SNF antagonism impairs tumor-induced expansion and suppressive function of PMN-MDSCs.** A. Gating strategy for mouse PMN-MDSCs. B. Left: Flow cytometric quantification of PMN-MDSC numbers in splenocytes isolated from 4T1 TB mice (left) or B16-F10 TB mice (middle) cultured in medium alone or supplemented with 25% TES. Right: Representative contour plots showing the proportional change of PMN-MDSCs treated in middle. C. Reverse transcription quantitative PCR assessing expression of *S100a8* and *S100a9* in TB PMN-MDSCs or TF neutrophils cultured in medium alone or supplemented with 25% TES, with 24-hour in vitro treatment by DMSO, FHD-286 (100 nM), or AU-24118 (3 µM). D. Gating strategy for human LOX-1^+^ PMN-MDSCs and LOX-1^-^ neutrophils. E. Assessment of PMN-MDSC migration following 24-hour pretreatment with DMSO, FHD_-_286 (100 nM), AU-15330 (3 µM), or AU-24118 (3 µM). F. Representative COMET™ multiplex immunofluorescence images showing S100A9 and CD8 expression in 4T1 mammary fat pad tumors following treatment with vehicle, AU_-_24118, or FHD-286. Scale bar, 20 µm. Right: Quantification shows the nearest neighbor distance between S100A9^+^ cells and CD8^+^ T cells across treatment groups. Data are presented as box and whisker plots in B, C, E and F, and were statistically assessed using two-way analysis of variance. Bonferroni’s correction is applied for multiple comparisons in F. All data are representative of two independent experiments.

**Figure S8. mSWI/SNF antagonism collapses PU.1- and C/EBPβ-dependent chromatin accessibility in PMN-MDSCs.** A. Venn diagrams showing overlaps of ATAC-seq peaks in DMSO, FHD-286 (100 nM), and AU-24118 (3 µM) treated PMN-MDSCs. B. ChIP-seq read-density heatmaps representing H3K27Ac at mSWI/SNF antagonists-loss genomic sites in PMN-MDSCs following 24 hour-treatment with DMSO, FHD-286 (100 nM), or AU-24118 (3 µM). C. Top: Top ten known motifs (ranked by *p*-value) enriched within C/EBPβ binding sites in PMN-MDSCs. Bottom: Top five known motifs (ranked by *p*-value) enriched within PU.1 binding sites in PMN-MDSCs.

## References

1. Lyu, A., et al., Evolution of myeloid-mediated immunotherapy resistance in prostate cancer. Nature, 2025. 637(8048).

2. Engblom, C., C. Pfirschke, and M.J. Pittet, The role of myeloid cells in cancer therapies. Nature Reviews Cancer, 2016. 16(7): p. 447–462.

3. Haynes, N.M., T.B. Chadwick, and B.S. Parker, The complexity of immune evasion mechanisms throughout the metastatic cascade. Nature Immunology, 2024. 25(10): p. 1793–1808.

4. Veglia, F., E. Sanseviero, and D.I. Gabrilovich, Myeloid-derived suppressor cells in the era of increasing myeloid cell diversity. Nat Rev Immunol, 2021. 21(8): p. 485–498.

5. Kumar, V., et al., The Nature of Myeloid-Derived Suppressor Cells in the Tumor Microenvironment. Trends in Immunology, 2016. 37(3): p. 208–220.

6. Passaro, A., et al., Gr-MDSC-linked asset as a potential immune biomarker in pretreated NSCLC receiving nivolumab as second-line therapy. Clinical & Translational Oncology, 2020. 22(4): p. 603–611.

7. Petrova, V., et al., Immunosuppressive capacity of circulating MDSC predicts response to immune checkpoint inhibitors in melanoma patients. Frontiers in Immunology, 2023. 14.

8. Sternberg, C., et al., *Randomized, Double-Blind,* Placebo-Controlled Phase III Study of Tasquinimod in Men With Metastatic Castration-Resistant Prostate Cancer. Journal of Clinical Oncology, 2016. 34(22): p. 2636–U136.

9. Greene, S., et al., Inhibition of MDSC Trafficking with SX-682, a CXCR1/2 Inhibitor, Enhances NK-Cell Immunotherapy in Head and Neck Cancer Models. Clinical Cancer Research, 2020. 26(6): p. 1420–1431.

10. Ho, L. and G.R. Crabtree, Chromatin remodelling during development. Nature, 2010. 463(7280): p. 474-84.

11. Kadoch, C., et al., Proteomic and bioinformatic analysis of mammalian SWI/SNF complexes identifies extensive roles in human malignancy. Nature Genetics, 2013. 45(6): p. 592-+.

12. Nakayama, R.T., et al., SMARCB1 is required for widespread BAF complex_-_mediated activation of enhancers and bivalent promoters. Nature Genetics, 2017. 49(11): p. 1613-+.

13. Xiao, L.B., et al., Targeting SWI/SNF ATPases in enhancer-addicted prostate cancer *(vol* 601, *pg* 434, 2022). Nature, 2024. 629(8011): p. E9-E9.

14. Malone, H.A. and C.W.M. Roberts, Chromatin remodellers as therapeutic targets. Nature Reviews Drug Discovery, 2024. 23(9): p. 661–681.

15. He, T.C., et al., Targeting the mSWI/SNF complex in POU2F-POU2AF transcription factor-driven malignancies. Cancer Cell, 2024. 42(8).

16. Liao, J.W., et al., Collaboration between distinct SWI/SNF chromatin remodeling complexes directs enhancer selection and activation of macrophage inflammatory genes. Immunity, 2024. 57(8).

17. Guo, A., et al., cBAF complex components and MYC cooperate early in CD8^+^ T cell fate. Nature, 2022. 607(7917): p. 135-+.

18. Baxter, A.E., et al., The SWI/SNF chromatin remodeling complexes BAF and PBAF differentially regulate epigenetic transitions in exhausted CD8+T cells. Immunity, 2023. 56(6): p. 1320-+.

19. He, T.C., et al., Development of an orally bioavailable mSWI/SNF ATPase degrader and acquired mechanisms of resistance in prostate cancer. Proceedings of the National Academy of Sciences of the United States of America, 2024. 121(15).

20. Vaswani, R.G., et al., Discovery of FHD-286, a First-in-Class, Orally Bioavailable, Allosteric Dual Inhibitor of the Brahma Homologue (BRM) and Brahma-Related Gene 1 (BRG1) ATPase Activity for the Treatment of SWItch/Sucrose Non_-_Fermentable (SWI/SNF) Dependent Cancers. Journal of Medicinal Chemistry, 2025. 68(2): p. 1772–1792.

21. He, T., et al., Development of an orally bioavailable mSWI/SNF ATPase degrader and acquired mechanisms of resistance in prostate cancer. Proc Natl Acad Sci U S A, 2024. 121(15): p. e2322563121.

22. He, T.C., et al., Targeting the mSWI/SNF complex in POU2F-POU2AF transcription factor-driven malignancies. Cancer Cell, 2024. 42(8).

23. Vaswani, R.G., et al., Discovery of FHD-286, a First-in-Class, Orally Bioavailable, Allosteric Dual Inhibitor of the Brahma Homologue (BRM) and Brahma-Related Gene 1 (BRG1) ATPase Activity for the Treatment of SWItch/Sucrose Non_-_Fermentable (SWI/SNF) Dependent Cancers. J Med Chem, 2025. 68(2): p. 1772–1792.

24. Wang, G.C., et al., Targeting YAP-Dependent MDSC Infiltration Impairs Tumor Progression. Cancer Discovery, 2016. 6(1): p. 80–95.

25. Bao, Y., et al., Targeting m6A reader YTHDF1 augments antitumour immunity and boosts anti-PD-1 efficacy in colorectal cancer. Gut, 2023. 72(8): p. 1497_-_1509.

26. Boivin, G., et al., Durable and controlled depletion of neutrophils in mice. Nature Communications, 2020. 11(1).

27. Kim, R., et al., Ferroptosis of tumour neutrophils causes immune suppression in cancer. Nature, 2022. 612(7939): p. 338-+.

28. SumiIchinose, C., et al., SNF2 beta-BRG1 is essential for the viability of F9 murine embryonal carcinoma cells. Molecular and Cellular Biology, 1997. 17(10): p. 5976–5986.

29. Veglia, F., et al., Fatty acid transport protein 2 reprograms neutrophils in cancer. Nature, 2019. 569(7754): p. 73-+.

30. Wang, C.X., et al., CD300ld on neutrophils is required for tumour-driven immune suppression. Nature, 2023. 621(7980): p. 830-+.

31. Reyes, J.C., et al., Altered control of cellular proliferation in the absence of mammalian brahma (SNF2alpha). EMBO J, 1998. 17(23): p. 6979–91.

32. Bultman, S., et al., A Brg1 null mutation in the mouse reveals functional differences among mammalian SWI/SNF complexes. Mol Cell, 2000. 6(6): p. 1287–95.

33. Shen, L., et al., Tasquinimod Modulates Suppressive Myeloid Cells and Enhances Cancer Immunotherapies in Murine Models. Cancer Immunology Research, 2015. 3(2): p. 136–148.

34. Zilionis, R., et al., Single-Cell Transcriptomics of Human and Mouse Lung Cancers Reveals Conserved Myeloid Populations across Individuals and Species. Immunity, 2019. 50(5): p. 1317-+.

35. Fang, J., et al., Genome-wide mapping of cancer dependency genes and genetic modifiers of chemotherapy in high-risk hepatoblastoma. Nature Communications, 2023. 14(1).

36. Hirai, H., et al., *C/EBP*β *is required for ‘emergency’ granulopoiesis*. Nature Immunology, 2006. 7(7): p. 732–739.

37. Marigo, I., et al., *Tumor-Induced Tolerance and Immune Suppression Depend on the C/EBP*β *Transcription Factor*. Immunity, 2010. 32(6): p. 790–802.

38. Strauss, L., et al., RORC1 Regulates Tumor-Promoting "Emergency" Granulo_-_Monocytopoiesis. Cancer Cell, 2015. 28(2): p. 253–269.

39. Gabrilovich, D.I. and S. Nagaraj, Myeloid-derived suppressor cells as regulators of the immune system. Nature Reviews Immunology, 2009. 9(3): p. 162–174.

40. Veglia, F., et al., Analysis of classical neutrophils and polymorphonuclear myeloid-derived suppressor cells in cancer patients and tumor-bearing mice. Journal of Experimental Medicine, 2021. 218(4).

41. Marvel, D. and D.I. Gabrilovich, Myeloid-derived suppressor cells in the tumor microenvironment: expect the unexpected. Journal of Clinical Investigation, 2015. 125(9): p. 3356–3364.

42. Li, W., et al., Aerobic Glycolysis Controls Myeloid-Derived Suppressor Cells and Tumor Immunity via a Specific CEBPB Isoform in Triple-Negative Breast Cancer. Cell Metabolism, 2018. 28(1): p. 87-+.

43. Minderjahn, J., et al., Mechanisms governing the pioneering and redistribution capabilities of the non-classical pioneer PU.1 *(vol 11*, *402*, *2020).* Nature Communications, 2020. 11(1).

44. Chambers, C., et al., SWI/SNF Blockade Disrupts PU.1-Directed Enhancer Programs in Normal Hematopoietic Cells and Acute Myeloid Leukemia. Cancer Research, 2023. 83(7): p. 983–996.

45. Kowenz-Leutz, E. and A. Leutz, *A C/EBP*β *isoform recruits the SWI/SNF complex to activate myeloid genes*. Molecular Cell, 1999. 4(5): p. 735–743.

46. Heinz, S., et al., Simple Combinations of Lineage-Determining Transcription Factors Prime cis-Regulatory Elements Required for Macrophage and B Cell Identities. Molecular Cell, 2010. 38(4): p. 576–589.

47. Guo, A., et al., cBAF complex components and MYC cooperate early in CD8^+^ T cell fate. Nature, 2022. 607(7917): p. 135–141.

48. Hegde, S., A.M. Leader, and M. Merad, MDSC: Markers, development, states, and unaddressed complexity. Immunity, 2021. 54(5): p. 875–884.

49. Lasser, S.A., et al., Myeloid-derived suppressor cells in cancer and cancer therapy. Nat Rev Clin Oncol, 2024. 21(2): p. 147–164.

50. Wang, P.F., et al., Prognostic role of pretreatment circulating MDSCs in patients with solid malignancies: A meta-analysis of 40 studies. Oncoimmunology, 2018. 7(10).

51. Weide, B., et al., Myeloid-Derived Suppressor Cells Predict Survival of Patients with Advanced Melanoma: Comparison with Regulatory T Cells and NY-ESO-1- or Melan-A-Specific T Cells. Clinical Cancer Research, 2014. 20(6): p. 1601_-_1609.

52. Guo, C., et al., Targeting myeloid chemotaxis to reverse prostate cancer therapy resistance. Nature, 2023. 623(7989): p. 1053-+.

53. Gatchalian, J., et al., Control of Stimulus-Dependent Responses in Macrophages by SWI/SNF Chromatin Remodeling Complexes. Trends in Immunology, 2020. 41(2): p. 126–140.

54. Ramirez-Carrozzi, V.R., et al., *Selective and antagonistic functions of SWI/SNF and Mi-2*β *nucleosome remodeling complexes during an inflammatory response*. Genes & Development, 2006. 20(3): p. 282–296.

55. Dinardo, C.D., et al., A Phase I Study of FHD-286, a Dual BRG1/BRM (SMARCA4/SMARCA2) Inhibitor, in Patients with Advanced Myeloid Malignancies. Clinical Cancer Research, 2025. 31(12): p. 2327–2338.

56. Bao, Y., et al., The UBA1-STUB1 Axis Mediates Cancer Immune Escape and Resistance to Checkpoint Blockade. Cancer Discovery, 2025. 15(2): p. 363–381.

57. Zheng, G.X.Y., et al., Massively parallel digital transcriptional profiling of single cells. Nature Communications, 2017. 8.

58. Young, M.D. and S. Behjati, SoupX removes ambient RNA contamination from droplet-based single-cell RNA sequencing data. Gigascience, 2020. 9(12).

59. Hao, Y., et al., Integrated analysis of multimodal single-cell data. Cell, 2021. 184(13): p. 3573–3587 e29.

60. Zhang, C.G., et al., Single-cell sequencing reveals antitumor characteristics of intratumoral immune cells in old mice. Journal for Immunotherapy of Cancer, 2021. 9(10).

61. Cable, D.M., et al., Robust decomposition of cell type mixtures in spatial transcriptomics. Nature Biotechnology, 2022. 40(4): p. 517-+.

62. Tien, J.C., et al., Defining CDK12 as a tumor suppressor and therapeutic target in mouse models of tubo-ovarian high-grade serous carcinoma. Proc Natl Acad Sci U S A, 2025. 122(24): p. e2426909122.

63. Ritchie, M.E., et al., *powers differential expression analyses for RNA-sequencing and microarray studies*. Nucleic Acids Research, 2015. 43(7).

64. Phipson, B., et al., Robust Hyperparameter Estimation Protects against Hypervariable Genes and Improves Power to Detect Differential Expression. Annals of Applied Statistics, 2016. 10(2): p. 946–963.

65. Robinson, M.D., D.J. McCarthy, and G.K. Smyth, edgeR: a Bioconductor package for differential expression analysis of digital gene expression data. Bioinformatics, 2010. 26(1): p. 139–140.

66. Wu, T.Z., et al., clusterProfiler 4.0: A universal enrichment tool for interpreting omics data. Innovation, 2021. 2(3).

67. Parolia, A., et al., NSD2 is a requisite subunit of the AR/FOXA1 neo_-_enhanceosome in promoting prostate tumorigenesis. Nature Genetics, 2024. 56(10).

68. Bolger, A.M., M. Lohse, and B. Usadel, Trimmomatic: a flexible trimmer for Illumina sequence data. Bioinformatics, 2014. 30(15): p. 2114–2120.

69. Li, H. and R. Durbin, Fast and accurate short read alignment with Burrows_-_Wheeler transform. Bioinformatics, 2009. 25(14): p. 1754–1760.

70. Danecek, P., et al., Twelve years of SAMtools and BCFtools. Gigascience, 2021. 10(2).

71. Institute, B. Picard Toolkit. Available from: http://broadinstitute.github.io/picard/.

72. Zhang, Y., et al., Model-based Analysis of ChIP-Seq (MACS). Genome Biology, 2008. 9(9).

73. Kent, W.J., et al., BigWig and BigBed: enabling browsing of large distributed datasets. Bioinformatics, 2010. 26(17): p. 2204–2207.

74. Yu, G.C., L.G. Wang, and Q.Y. He, ChIPseeker: an R/Bioconductor package for ChIP peak annotation, comparison and visualization. Bioinformatics, 2015. 31(14): p. 2382–2383.

75. Zhu, L.J., Integrative Analysis of ChIP-Chip and ChIP-Seq Dataset. Tiling Arrays: Methods and Protocols, 2013. 1067: p. 105–124.

76. Wang, Q.W., et al., Exploring Epigenomic Datasets by ChIPseeker. Current Protocols, 2022. 2(10).

77. Ramírez, F., et al., deepTools2: a next generation web server for deep_-_sequencing data analysis. Nucleic Acids Research, 2016. 44(W1): p. W160_-_W165.

