## Supplemental Figures S1 - S8 for "A druggable chromatin vulnerability in suppressive neutrophils restores antitumor immunity"

### Figure S1

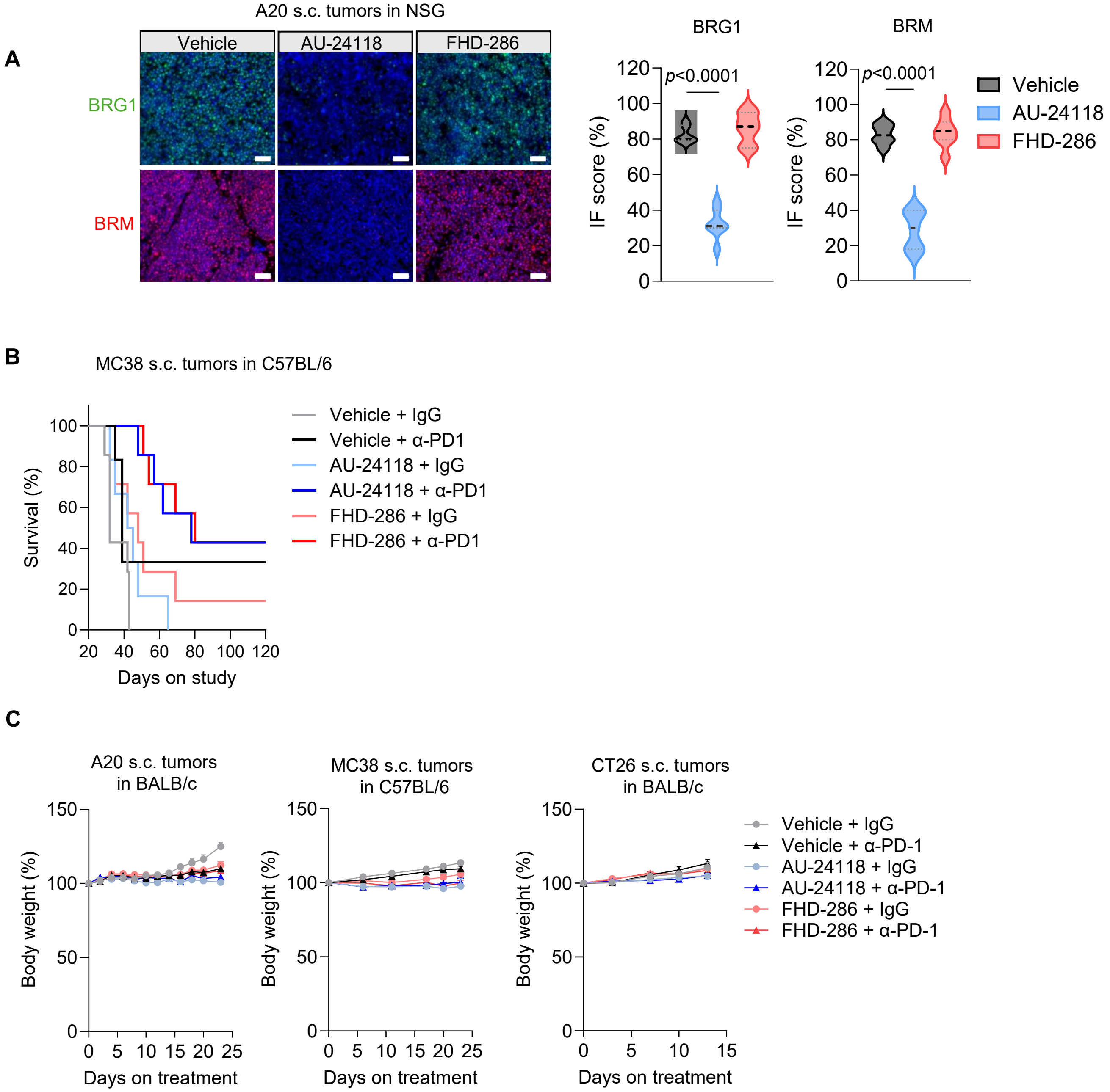
A
A20 s.c. tumors in NSG
Vehicle
AU-24118
FHD-286
BRG1
BRM
BRG1
BRM
IF score (%)
p<0.0001
Vehicle
AU-24118
FHD-286
B
MC38 s.c. tumors in C57BL/6
Survival (%)
Days on study
Vehicle + IgG
Vehicle + α-PD1
AU-24118 + IgG
AU-24118 + α-PD1
FHD-286 + IgG
FHD-286 + α-PD1
C
A20 s.c. tumors in BALB/c
MC38 s.c. tumors in C57BL/6
CT26 s.c. tumors in BALB/c
Body weight (%)
Days on treatment
Vehicle + IgG
Vehicle + α-PD-1
AU-24118 + IgG
AU-24118 + α-PD-1
FHD-286 + IgG
FHD-286 + α-PD-1

#### Figure S2

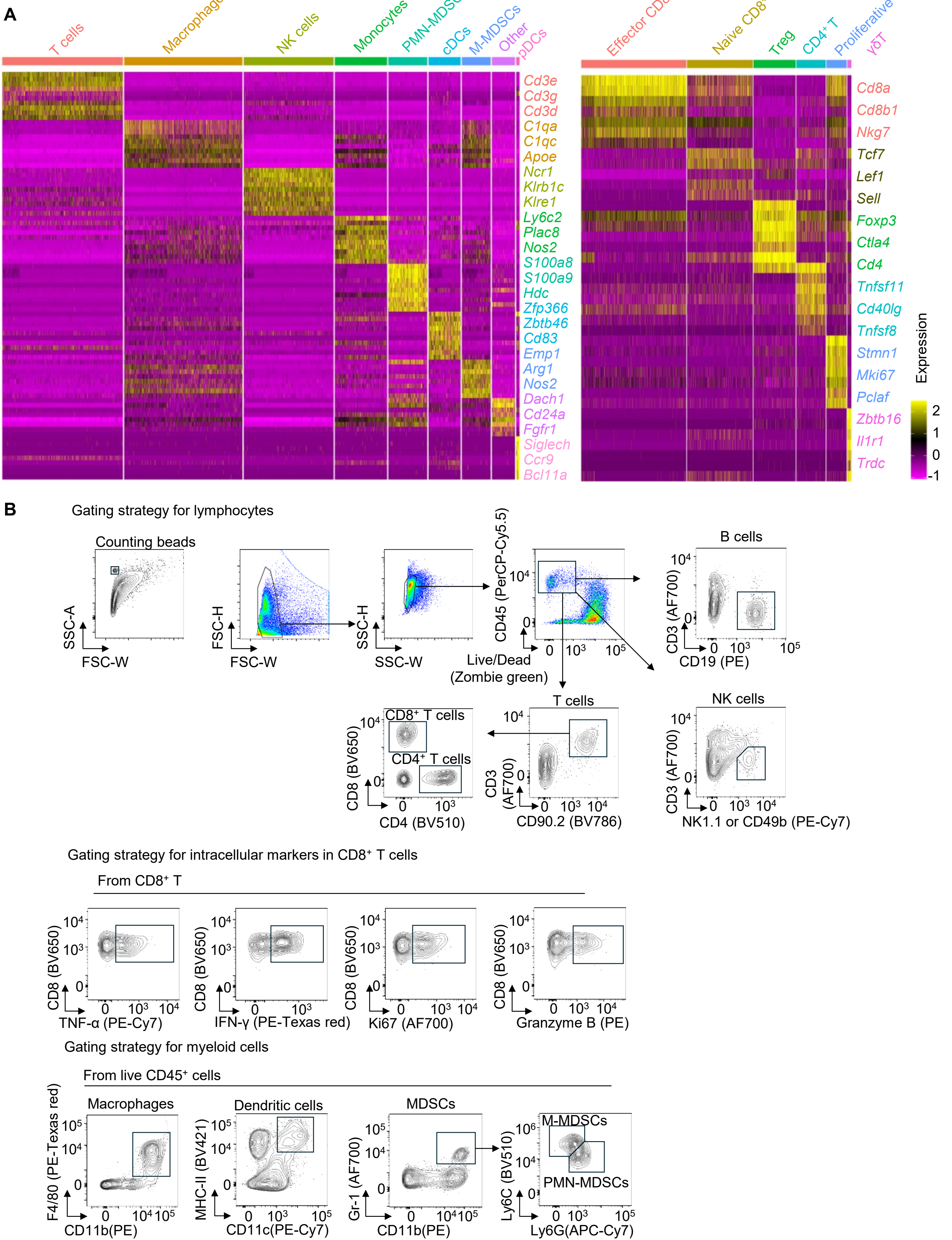
A
T cells
Macrophage
NK cells
Monocytes
PMN-MDSCs
cDCs
M-MDSCs
Other
pDCs
Cd3e
Cd3g
Cd3d
C1qa
C1qc
Apoe
Ncr1
Klrb1c
Klre1
Ly6c2
Plac8
Nos2
S100a8
S100a9
Hdc
Zfp366
Zbtb46
Cd83
Emp1
Arg1
Nos2
Dach1
Cd24a
Fgfr1
Siglech
Ccr9
Bcl11a
Effector CD8+ T
Naive CD8+ T
Treg
CD4+ T
Proliferative CD8+ T
γδT
Cd8a
Cd8b1
Nkg7
Tcf7
Lef1
Sell
Foxp3
Ctla4
Cd4
Tnfsf11
Cd40lg
Tnfsf8
Stmn1
Mki67
Pclaf
Zbtb16
Il1r1
Trdc
Expression
2
1
0
-1
B
Gating strategy for lymphocytes
Counting beads
SSC-A
FSC-W
FSC-H
FSC-W
SSC-H
SSC-W
CD45 (PerCP-Cy5.5)
Live/Dead (Zombie green)
B cells
CD3 (AF700)
CD19 (PE)
T cells
CD3 (AF700)
CD90.2 (BV786)
CD8+ T cells
CD4+ T cells
CD8 (BV650)
CD4 (BV510)
NK cells
CD3 (AF700)
NK1.1 or CD49b (PE-Cy7)
Gating strategy for intracellular markers in CD8+ T cells
From CD8+ T
CD8 (BV650)
TNF-α (PE-Cy7)
IFN-γ (PE-Texas red)
Ki67 (AF700)
Granzyme B (PE)
Gating strategy for myeloid cells
From live CD45+ cells
Macrophages
F4/80 (PE-Texas red)
CD11b(PE)
Dendritic cells
MHC-II (BV421)
CD11c(PE-Cy7)
MDSCs
Gr-1 (AF700)
CD11b(PE)
M-MDSCs
PMN-MDSCs
Ly6C (BV510)
Ly6G(APC-Cy7)

### Figure S3

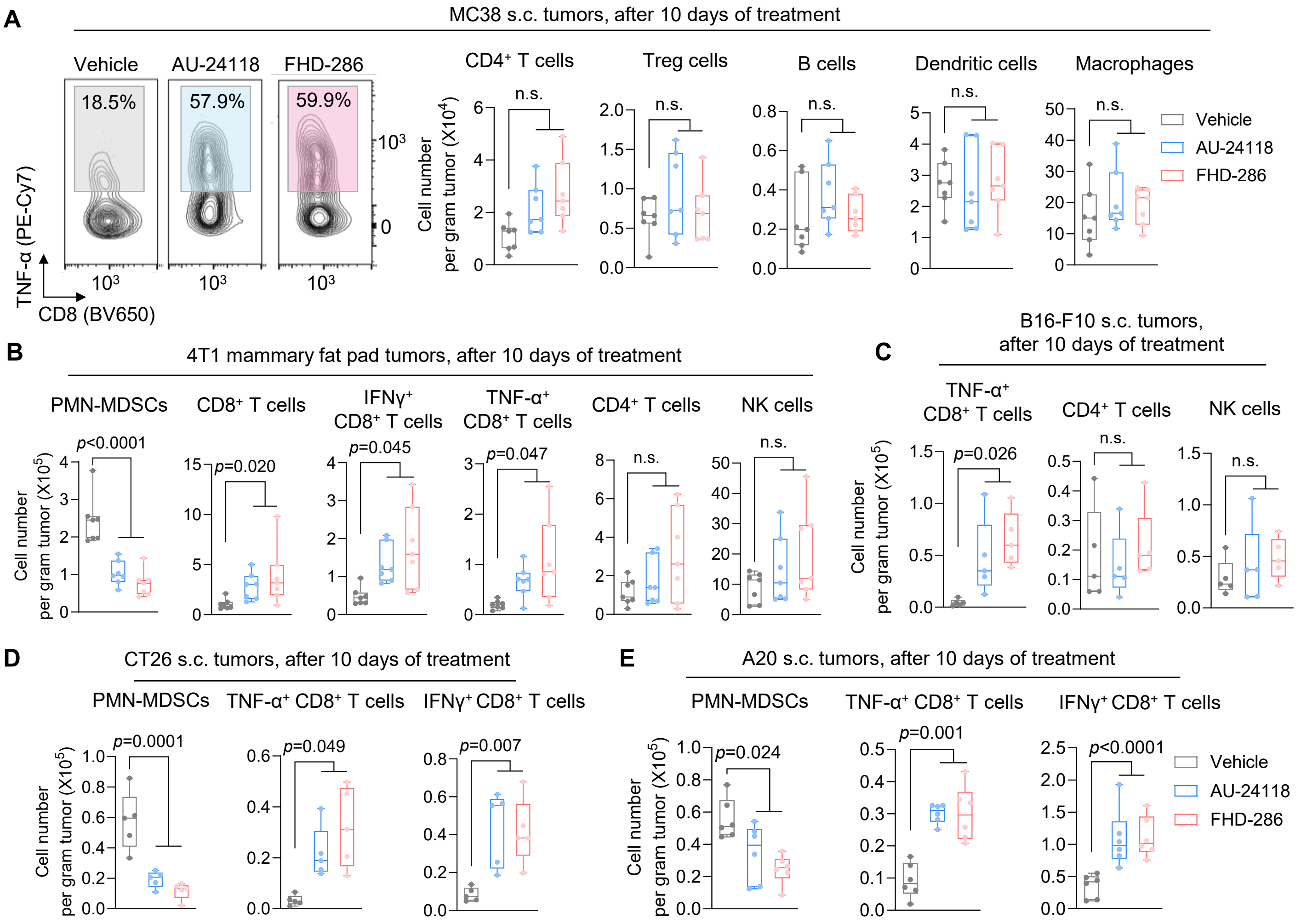

A
MC38 s.c. tumors, after 10 days of treatment
Vehicle
AU-24118
FHD-286
18.5%
57.9%
59.9%
TNF-α (PE-Cy7)
CD8 (BV650)
CD4⁺ T cells
Treg cells
B cells
Dendritic cells
Macrophages
n.s.
Cell number per gram tumor (X10⁴)
B
4T1 mammary fat pad tumors, after 10 days of treatment
PMN-MDSCs
CD8⁺ T cells
IFNγ⁺ CD8⁺ T cells
TNF-α⁺ CD8⁺ T cells
CD4⁺ T cells
NK cells
p<0.0001
p=0.020
p=0.045
p=0.047
Cell number per gram tumor (X10⁵)
C
B16-F10 s.c. tumors, after 10 days of treatment
TNF-α⁺ CD8⁺ T cells
CD4⁺ T cells
NK cells
p=0.026
D
CT26 s.c. tumors, after 10 days of treatment
PMN-MDSCs
TNF-α⁺ CD8⁺ T cells
IFNγ⁺ CD8⁺ T cells
p=0.0001
p=0.049
p=0.007
E
A20 s.c. tumors, after 10 days of treatment
PMN-MDSCs
TNF-α⁺ CD8⁺ T cells
IFNγ⁺ CD8⁺ T cells
p=0.024
p=0.001
p<0.0001

#### Figure S4

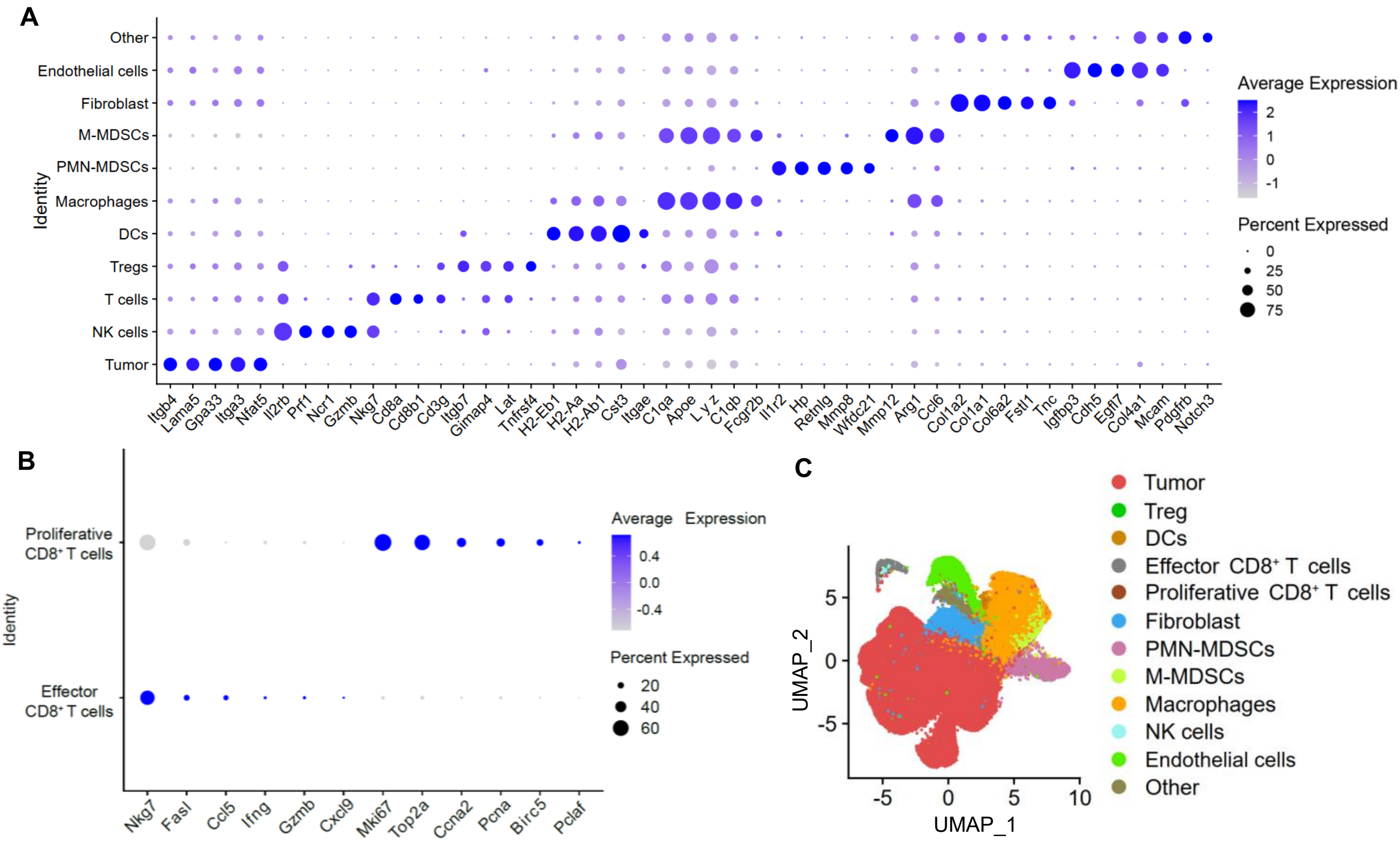
A
Other
Endothelial cells
Fibroblast
M-MDSCs
PMN-MDSCs
Macrophages
DCs
Tregs
T cells
NK cells
Tumor
Identity
Itgb4
Lama5
Gpa33
Itga3
Nfat5
Il2rb
Prf1
Ncr1
Gzmb
Nkg7
Cd8a
Cd8b1
Cd3g
Itgb7
Gimap4
Lat
Tnfrsf4
H2-Eb1
H2-Aa
H2-Ab1
Cst3
Itgae
C1qa
Apoe
Lyz
C1qb
Fcgr2b
Il1r2
Hp
Retnlg
Mmp8
Wfdc21
Mmp12
Arg1
Ccl6
Col1a2
Col1a1
Col6a2
Fstl1
Tnc
Igfbp3
Cdh5
Egfl7
Col4a1
Mcam
Pdgfrb
Notch3
Average Expression
2
1
0
-1
Percent Expressed
0
25
50
75
B
Proliferative CD8⁺ T cells
Effector CD8⁺ T cells
Identity
Nkg7
Fasl
Ccl5
Ifng
Gzmb
Cxcl9
Mki67
Top2a
Ccna2
Pcna
Birc5
Pclaf
Average Expression
0.4
0.0
-0.4
Percent Expressed
20
40
60
C
UMAP_2
UMAP_1
5
0
-5
-5
0
5
10
Tumor
Treg
DCs
Effector CD8⁺ T cells
Proliferative CD8⁺ T cells
Fibroblast
PMN-MDSCs
M-MDSCs
Macrophages
NK cells
Endothelial cells
Other

### Figure S5

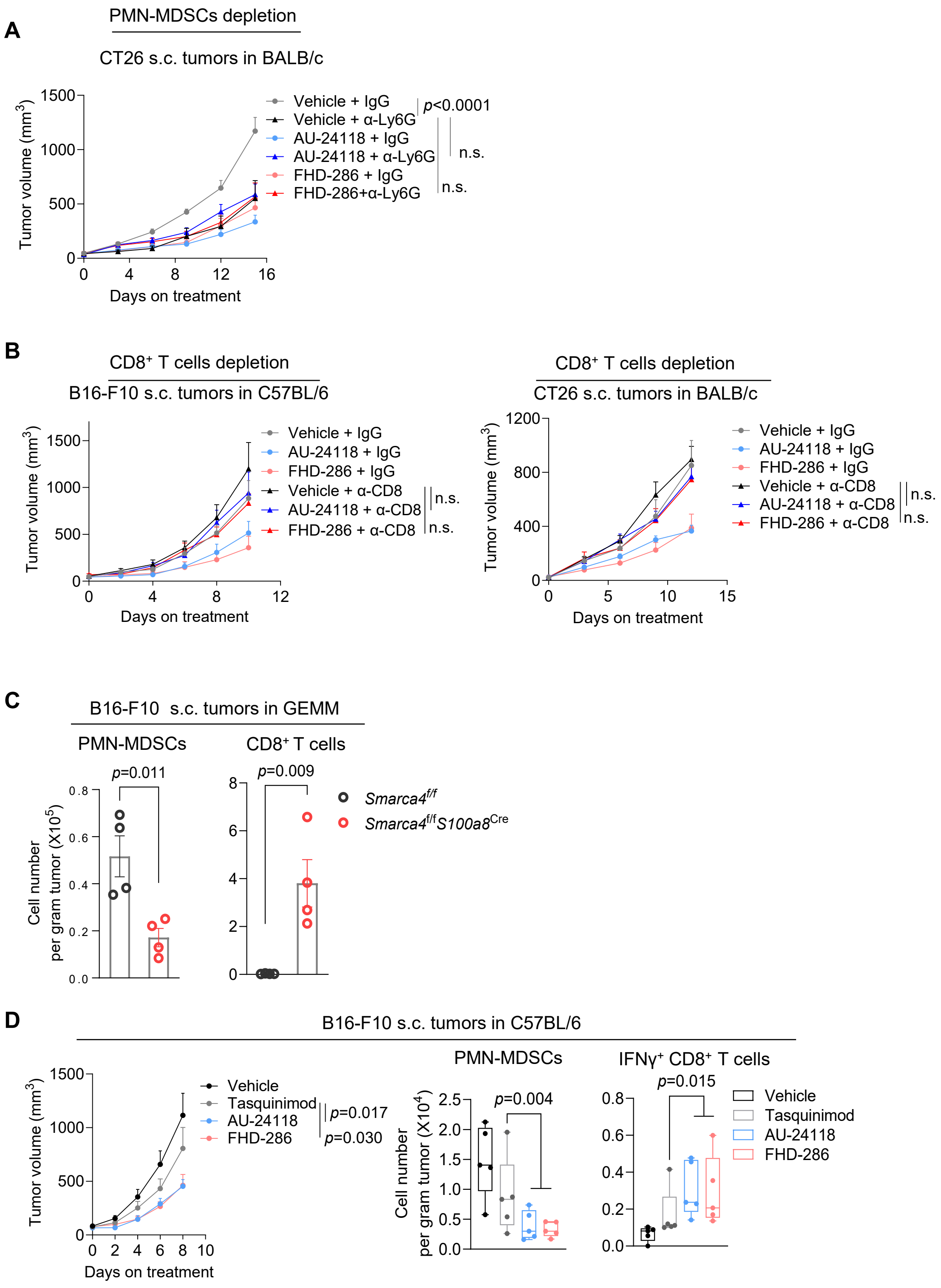

A
PMN-MDSCs depletion
CT26 s.c. tumors in BALB/c
Tumor volume (mm³)
1500
1000
500
0
0 4 8 12 16
Days on treatment
Vehicle + IgG
Vehicle + α-Ly6G
AU-24118 + IgG
AU-24118 + α-Ly6G
FHD-286 + IgG
FHD-286+α-Ly6G
p<0.0001
n.s.
n.s.
B
CD8⁺ T cells depletion
B16-F10 s.c. tumors in C57BL/6
Tumor volume (mm³)
1500
1000
500
0
0 4 8 12
Days on treatment
Vehicle + IgG
AU-24118 + IgG
FHD-286 + IgG
Vehicle + α-CD8
AU-24118 + α-CD8
FHD-286 + α-CD8
n.s.
n.s.
CD8⁺ T cells depletion
CT26 s.c. tumors in BALB/c
Tumor volume (mm³)
1200
800
400
0
0 5 10 15
Days on treatment
Vehicle + IgG
AU-24118 + IgG
FHD-286 + IgG
Vehicle + α-CD8
AU-24118 + α-CD8
FHD-286 + α-CD8
n.s.
n.s.
C
B16-F10 s.c. tumors in GEMM
PMN-MDSCs
CD8⁺ T cells
p=0.011
p=0.009
Cell number per gram tumor (X10⁵)
0.8
0.6
0.4
0.2
0.0
8
6
4
2
0
Smarca4f/f
Smarca4f/fS100a8Cre
D
B16-F10 s.c. tumors in C57BL/6
Tumor volume (mm³)
1500
1000
500
0
0 2 4 6 8 10
Days on treatment
Vehicle
Tasquinimod
AU-24118
FHD-286
p=0.017
p=0.030
PMN-MDSCs
Cell number per gram tumor (X10⁴)
2.5
2.0
1.5
1.0
0.5
0.0
p=0.004
IFNγ⁺ CD8⁺ T cells
0.8
0.6
0.4
0.2
0.0
p=0.015
Vehicle
Tasquinimod
AU-24118
FHD-286

### Figure S6

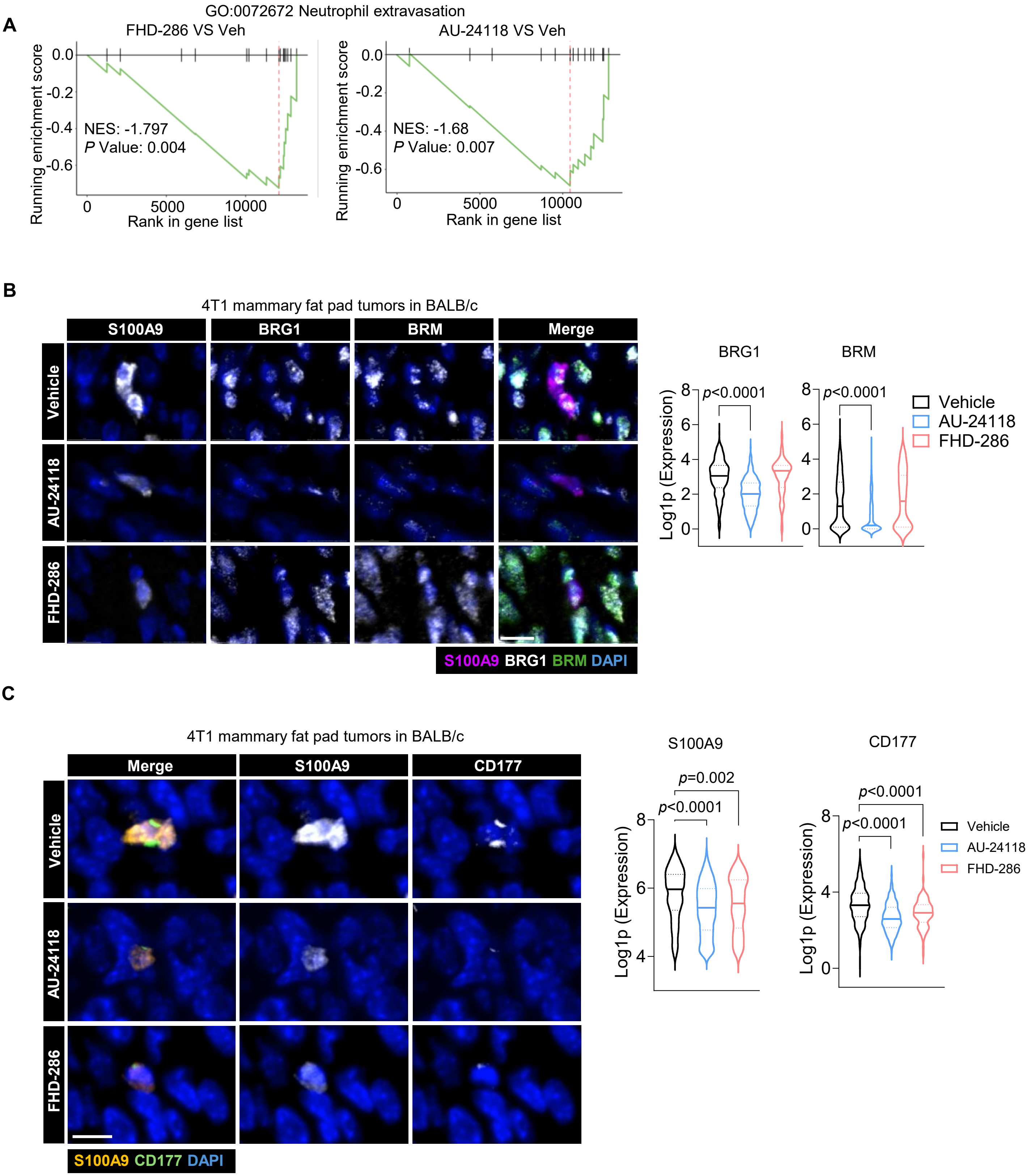
A
GO:0072672 Neutrophil extravasation
FHD-286 VS Veh
AU-24118 VS Veh
Running enrichment score
NES: -1.797
P Value: 0.004
NES: -1.68
P Value: 0.007
Rank in gene list
B
4T1 mammary fat pad tumors in BALB/c
S100A9
BRG1
BRM
Merge
Vehicle
AU-24118
FHD-286
S100A9 BRG1 BRM DAPI
BRG1
BRM
Log1p (Expression)
p<0.0001
p<0.0001
Vehicle
AU-24118
FHD-286
C
4T1 mammary fat pad tumors in BALB/c
Merge
S100A9
CD177
Vehicle
AU-24118
FHD-286
S100A9 CD177 DAPI
S100A9
CD177
Log1p (Expression)
p=0.002
p<0.0001
p<0.0001
p<0.0001
Vehicle
AU-24118
FHD-286

### Figure S7

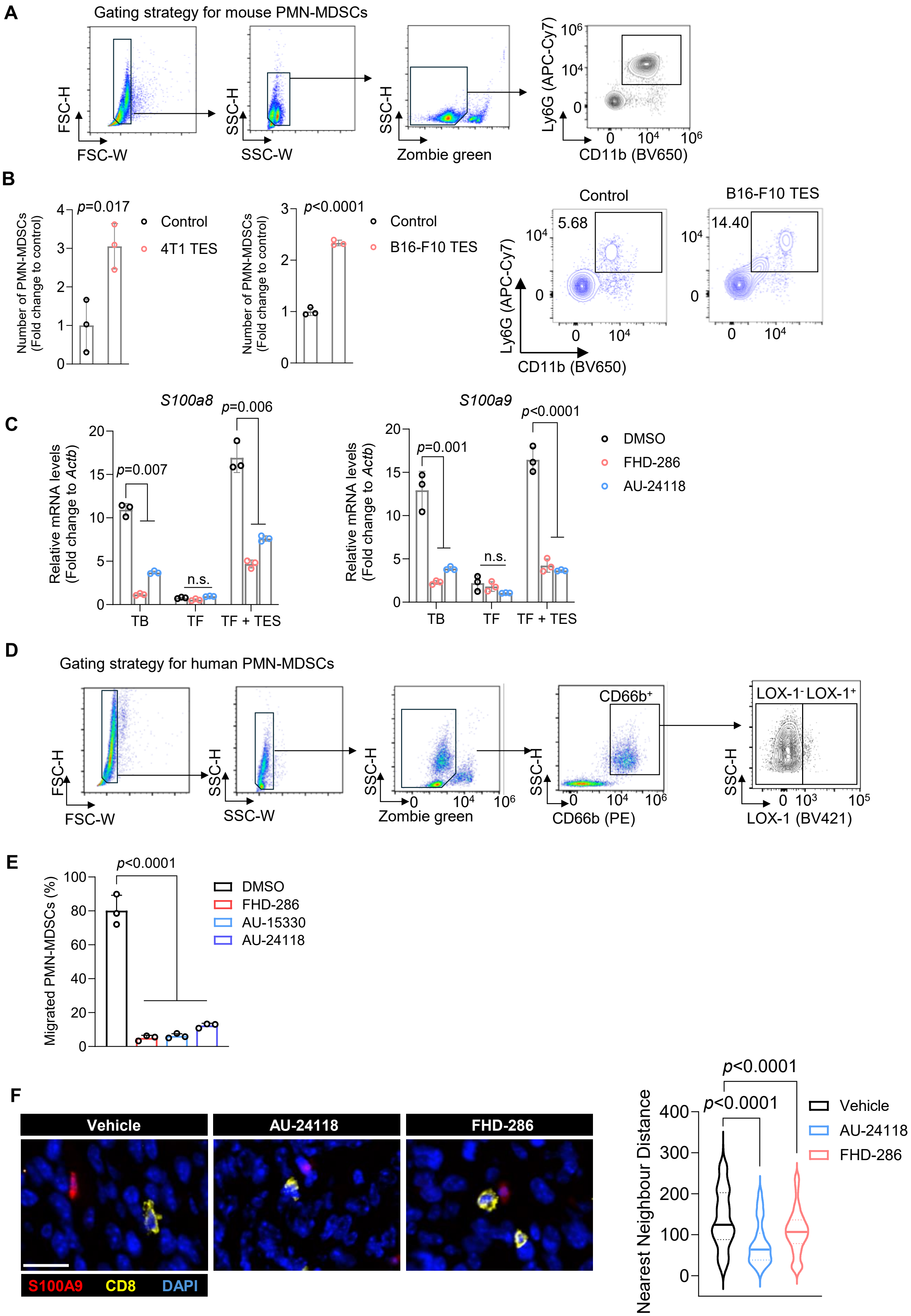

A
Gating strategy for mouse PMN-MDSCs
FSC-H
FSC-W
SSC-H
SSC-W
SSC-H
Zombie green
Ly6G (APC-Cy7)
CD11b (BV650)
B
Number of PMN-MDSCs (Fold change to control)
p=0.017
Control
4T1 TES
p<0.0001
Control
B16-F10 TES
Control
B16-F10 TES
5.68
14.40
Ly6G (APC-Cy7)
CD11b (BV650)
C
S100a8
S100a9
Relative mRNA levels (Fold change to Actb)
p=0.007
p=0.006
p=0.001
p<0.0001
n.s.
n.s.
TB
TF
TF + TES
DMSO
FHD-286
AU-24118
D
Gating strategy for human PMN-MDSCs
FSC-H
FSC-W
SSC-H
SSC-W
SSC-H
Zombie green
SSC-H
CD66b⁺
CD66b (PE)
SSC-H
LOX-1⁻ LOX-1⁺
LOX-1 (BV421)
E
Migrated PMN-MDSCs (%)
p<0.0001
DMSO
FHD-286
AU-15330
AU-24118
F
Vehicle
AU-24118
FHD-286
S100A9
CD8
DAPI
Nearest Neighbour Distance
p<0.0001
p<0.0001
Vehicle
AU-24118
FHD-286

**Figure S8**

**A**

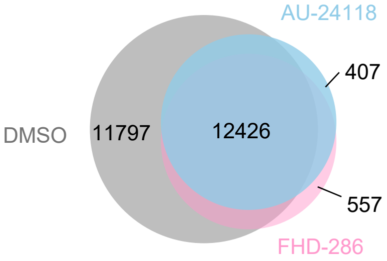


**B**

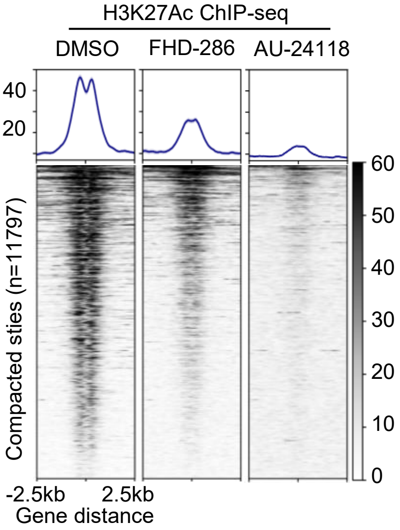


**C**

Homer Motif Enrichment (PMN-MDSCs C/EBPβ ChIP-seq, Top 10 motifs)

| Rank | Known motif | Name | log p-value | % Target with Motif | % Background with Motif |
|---|---|---|---|---|---|
| 1 | 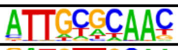 | CEBP | -7.984e+02 | 39.93% | 16.58% |
| 2 | 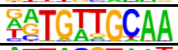 | CEBP:AP1 | -4.697e+02 | 35.95% | 18.02% |
| 3 | 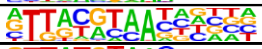 | NFIL3 | -3.493e+02 | 31.69% | 16.76% |
| 4 | 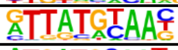 | HLF | -3.399e+02 | 38.08% | 22.10% |
| 5 | 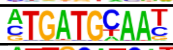 | Atf4 | -3.238e+02 | 16.45% | 6.31% |
| 6 | 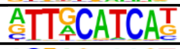 | Chop | -2.318e+02 | 12.45% | 4.87% |
| 7 | 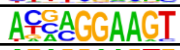 | ELF5 | -1.643e+02 | 27.45% | 17.39% |
| 8 | 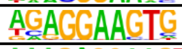 | PU.1 | -1.452e+02 | 19.43% | 11.36% |
| 9 | 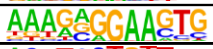 | SpiB | -1.409e+02 | 11.29% | 5.37% |
| 10 | 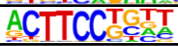 | Elf4 | -1.368e+02 | 32.13% | 22.26% |

Homer Motif Enrichment (PMN-MDSCs PU.1 ChIP-seq, Top 5 motifs)

| Rank | Known motif | Name | log p-value | % Target with Motif | % Background with Motif |
|---|---|---|---|---|---|
| 1 | 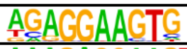 | PU.1 | -1.984e+04 | 51.86% | 11.64% |
| 2 | 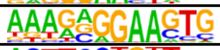 | SpiB | -1.973e+04 | 37.69% | 5.28% |
| 3 | 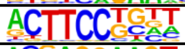 | Elf4 | -1.532e+04 | 65.84% | 24.83% |
| 4 | 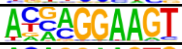 | ELF5 | -1.284e+04 | 51.97% | 17.23% |
| 5 | 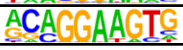 | ETS1 | -1.233e+04 | 61.32% | 24.71% |
